# Weakly deleterious natural genetic variation greatly amplifies probability of resistance in multiplexed gene drive systems

**DOI:** 10.1101/2021.12.23.473701

**Authors:** Bhavin S. Khatri, Austin Burt

## Abstract

Evolution of resistance is a major barrier to successful deployment of gene drive systems to suppress natural populations, which could greatly reduce the burden of many vector borne diseases. Multiplexed guide RNAs that require resistance mutations in all target cut sites is a promising anti-resistance strategy, since in principle resistance would only arise in unrealistically large populations. Using novel stochastic simulations that accurately model evolution at very large population sizes, we explore the probability of resistance due to three important mechanisms: 1) non-homologous end-joining mutations, 2) single nucleotide mutants arising de novo or, 3) single nucleotide polymorphisms pre-existing as standing variation. Our results explore the relative importance of these mechanisms and highlight a complexity of the mutation-selection-drift balance between haplotypes with complete resistance and those with an incomplete number of resistant alleles. We find this leads to a qualitatively new phenomenon where weakly deleterious naturally occurring variants greatly amplify the probability of multi-site resistance. This challenges the intuition that many target sites would guarantee prevention of resistance, where in the face of standing genetic variation, it can be probable even in not very large populations. This result has broad application to resistance arising in many multi-site evolutionary scenarios including multi-drug resistance to antibiotics, antivirals and cancer treatments, as well as the evolution of vaccine escape mutations in large populations.

## Introduction

Suppression gene drive systems have the potential to be highly effective for population control of many vectors of human disease, including malaria carrying mosquitoes, as well as for invasive species control (*1*). With the recent discovery of the CRISPR-Cas9 gene editing system (*2*), such drives can be realised by targeting specific sequences for insertion of a drive construct into a gene of importance; during meiosis this converts a very large fraction gametes carrying wild type alleles to this drive construct, thereby increasing transmission of drive over and above random Mendelian segregation. If the drive construct only has significant fitness effects as a homozygote, it can rise to high frequencies; in particular, if the target gene is necessary for female fertility this provides an effective way to suppress the population (*3, 4*). An important question is how resilient gene drive systems are to the evolution of resistance (*1*, *5–7*), which could arise via single nucleotide mutations or polymorphisms (SNPs) or by an imperfect end-joining repair process called non-homologous end joining (NHEJ) (*3, 8*). Although, in caged populations resistance to suppression has been observed via indels induced by NHEJ (*3*), drive systems where the chosen target site is critical to female fertility have been found to be resilient (*4*). However, it is an open question as to how resilient gene drive systems are to the evolution of resistance in natural populations, which can have very large population sizes (*9*).

Although empirically we understand well the rate at which these processes occur in individuals, from characterisation of the mutation rate (*µ*) and the net NHEJ rate (*ν*), both per generation, it is also crucially important to know 1) the population size, and 2) the fraction of potential resistance mutants that are actually functional. Rather than the per individual rate, the population-level mutation rate is critical, as it measures how likely such mutations are to arise in the population (*5–7*). For example, if a resistance mutant is produced per individual at rate 10^−7^ and the population size is *N* = 10^4^, the population level mutation rate is 10^−3^ and it will typically take 1000 generations before a single resistance mutant is generated in a population, and so resistance is unlikely to arise given that drive typically acts to suppress a population on a timescale of much less than 100 generations; on the other hand if *N* = 10^8^, 10 such mutants are generated every generation, and so resistance mutants with a selective advantage compared to drive are very likely to arise and fix before population elimination.

A promising anti-resistance strategy is to exploit multiple redundant target sites for cleavage by the nuclease (*1, 7, 10, 11*). If there are multiple guide RNAs (gRNAs) or target cut sites within a single locus, since a single successful cut, followed by homology directed repair, is sufficient for drive to be copied, all sites must develop resistance mutants in order for the non-Mendelian transmission of drive to fail. This aims to reduce the individual rate of resistance to such small numbers that typically unrealistic population sizes would be required to achieve a large population level rate of resistance generation. So although, this strategy has been demonstrated to work in cage trials (*12*), extrapolation of these results to large natural populations is uncertain. Using a deterministic model, Noble et al. (*11*), showed that multiplexed strategies can allow drive to overcome resistance on short time scales, however, as population dynamics were not considered, in such infinite population size models, resistance will always arise eventually for any fertile allele. Beaghton et al, explored explicit population dynamics and finite population stochasticity and the probability of resistance using branching process theory in the context of the Y-drive suppression strategy (*5*) and Marshall et al (*7*) used more complex and realistic stochastic simulations to examine how resistance interplays with population size and the rate of generation of NHEJ mutants. Both these works showed that large natural population sizes reduce the maximum permissible rate of resistance mutants. Using heuristic arguments, Marhsall et al, then argued that relatively modest degrees of multiplexing *m* could prevent resistance from arising. These works have not explored the role of standing variation, but more recently Lanzaro et al (*13*) showed that higher levels of standing variation of resistance variants increases the probability of resistance to various gene drive strategies including suppression drives, though crucially they did not look at the role of population size and multiplexing.

On a theoretical level there are a number of works in the population genetic literature of relevance to resistance in suppression drives (*14–17*) which predict that population rescue or adaptation in response to an environmental change, in general, is more likely from standing variation than compared to de novo mutants arising. Although simple theoretical arguments for the role of standing variation with multi-site evolution have been considered previously using branching process theory (*18*), they assumed the initial frequency of mutants are fixed at their deterministic mutation-balance frequencies. However, stochastic theory (*17*) shows that this overestimates the role of standing variation, since by averaging over the allele frequency distribution before the change, the advantage is only weakly logarithmic in the fitness cost, and in general not very significant. There is currently no theory that addresses the question of the role that standing variation plays, with its full allele frequency spectrum, in determining the probability of resistance for a multi-site evolutionary system like multiplexed gene-drive, and this is an important question we address here.

In this work, we develop fully stochastic Wright-Fisher simulations of multiplexed drive to analyse the role of population size in more detail, for different degrees of multiplex *m* and for more general scenarios of resistance not previously considered, including, as we shall see, the critical role of standing variation in multiplexed drives. We exclusively explore the population rescue scenario, where drive is introduced at sufficiently large frequency that the population will be eliminated without resistance. Multiplexing means the critical population sizes required for resistance can be very large, yet the resistance mutants still arise at very small frequency, requiring a stochastic description of their establishment; to this end, we develop a new and very accurate Gaussian-Poisson approximation to the multinomial distribution to allow Wright-Fisher sampling in almost arbitrarily large population sizes. We use these simulations to assess the modes of resistance arising for both NHEJ mutants and SNP mutants, where the later can arise by de novo base-pair mutation or pre-exist in the population before the introduction of drive as standing genetic variation. Importantly, we also allow each type to be functional or non-functional. Our key new finding shows the more complex structure of standing variation with multiple gRNAs leads to a power law dependence on the fitness cost of standing variation, compared to the weak logarithmic fitness effects for a single gRNA. This leads to an extreme sensitivity of resistance on the fitness cost, greatly amplifying the probability of resistance, compared to de novo, for weakly selected alleles as the number of target sites and gRNAs increases. As well as being important to predict the success of multiplexed drives systems in preventing resistance at a given population size, we suggest this result has wide applicability to understanding resistance due to standing variation for a range of multi-site evolutionary systems, including the question of vaccine escape.

## Results

Our main results are the probability of resistance arising as a function of the effective population size *N*, for one, two, or three gRNAs. We will explore the probability of resistance as a function of three main parameters, the fraction of NHEJ mutants that are functional (*β*), the fraction of de novo mutations that are functional (*ξ*), and the heterozygous fitness cost (in the absence of drive) of functional resistance (*σ*) (assumed to be the same whether it was derived from de novo mutation or NHEJ); the remaining fraction 1 − *β* of NHEJ mutants, or 1 − *ξ* of de novo SNPs are assumed nonfunctional. We find that the probability of resistance has a universal sigmoidal behaviour, which is of the form

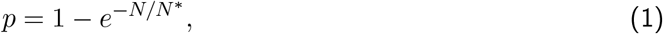

where for small population sizes *N* ≪ *N*^∗^ resistance is very unlikely, whilst for sufficiently large population sizes (*N* ≫ *N*^∗^) resistance arises with near 100% certainty. We use the simulations to develop a heuristic theory to characterise how *N*^∗^ depends on the parameters *β, ξ, σ, N* and then focus in particular on contours in the parameter space of these variables where the probability of resistance is *p* = 0.05, or 5%.

The results are organised by 4 different conditions: 1) NHEJ mutations only, 2) de novo SNP mutants only, 3) standing variation of SNP mutants in mutation-selection balance, and 4) where all of the first three mechanisms are possible, where for each we vary the fraction of functional NHEJ mutants *β*, the fraction of de novo SNPs *ξ*, and the fitness costs of functional mutants (NHEJ or SNP) as appropriate.

All the results are determined from running 500 replicate simulations. For each set of parameters we determine whether resistance to the drive construct arose using a resistance criterion that the sum of the frequency of resistance allele/haplotypes reaches 0.95; for example, when the frequency of R and N, for a single gRNA, is greater than 95%, or for 2 gRNAs, when the frequency of RR, RN and NN is greater than 90%. We also examine the time to resistance, given that resistance arises, in the Supplementary material. In all simulations drive is introduced at a frequency of 0.1 in males.

### Probability of resistance: NHEJ only and single nucleotide de novo mutants only

For both NHEJ and de novo single nucleotide mutants, we find that the simulations of the probability of resistance as a function of population can be fit very accurately with a sigmoidal form *p* = 1 − *e*^−*N/N*∗^. For NHEJ only we plot in Fig.1 *p* vs *N*, for a) *m* = 1 gRNA, b) *m* = 2 gRNAs and c) *m* = 3 gRNAs, for the standard parameters outlined above, but where the cost of functional resistance mutants is *σ* = 0.01. Each set of simulation data with square symbols of a given colour corresponds to a different value of *β*, the fraction of functional NHEJ mutants; as *β* decreases *N*^∗^ increases, as we would expect and the probability of resistance decreases for a given *N*. We find equivalent sigmoidal curves for de novo SNPs only, where *N*^∗^ increases for decreasing *ξ*. Again for NHEJ only, in Supplementary Information we show typical time series of the allele frequencies and population dynamics, showing population extinction for *N* < *N*^∗^ and population recovery and resistance for *N* > *N*^∗^, when the resistance allele/haplotype R, RR or RRR fix in the population, for *m* = 1, *m* = 2 and *m* = 3, respectively (Fig.S3). Note these results (square symbols) are obtained using the hybrid Poisson-Gaussian approximation to the multinomial distribution (see Methods), which allows Wright-Fisher simulations at very large population sizes; to check this approximation works well, the results with pentagram symbols represent simulations at smaller population sizes using multinomial generated random numbers, for which we see very good agreement with the Poisson-Gaussian approximation.

**Figure 1:**
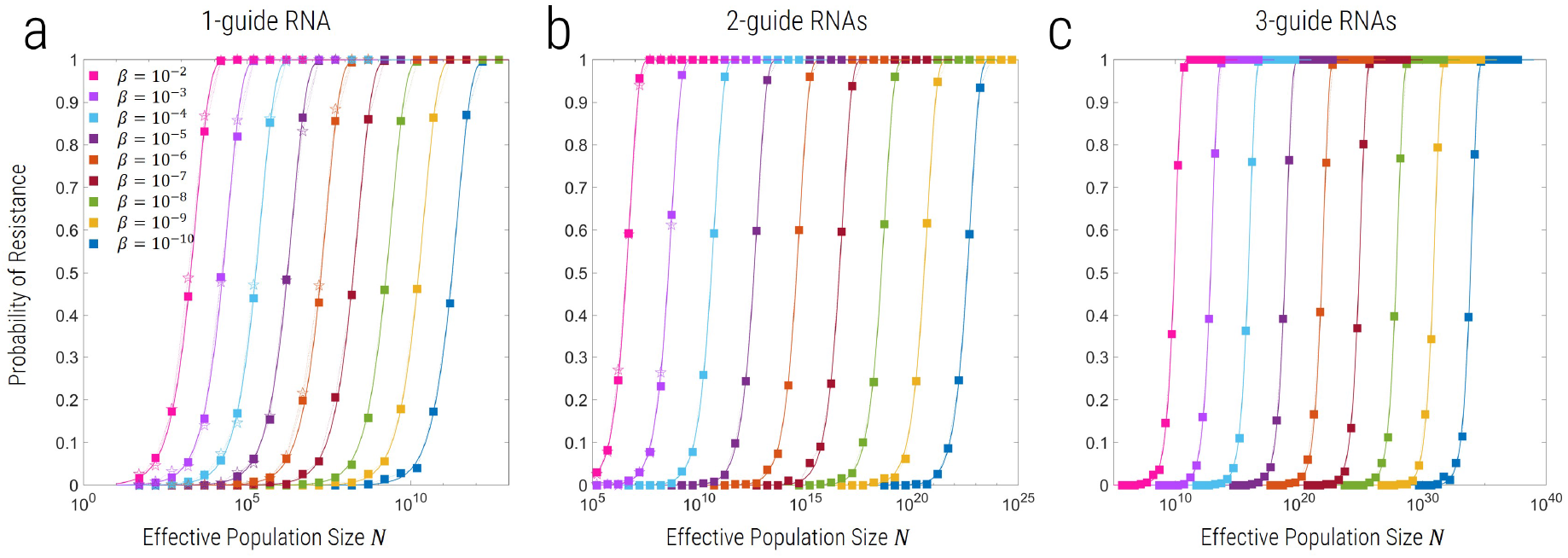
Probability of resistance evolving as a function of effective population size (*N*) for *m* = 1, 2, or 3 gRNAs, where resistance can only arise due to functional NHEJ mutants. R alleles have a fitness cost of *σ* = 0.01 relative to the wild type. Curves of different colour correspond to different values of *β*, where squares correspond to simulations using the hybrid Poisson-Gaussian approximation and open pentagrams to simulations using exact multinomial sampling.

The dependence of *N*^∗^ on *β* and *ξ* for NEHJ and de novo SNPs, respectively are similar, but with a crucial difference that highlights a difference in the dynamics of multiplex allele generation. For NHEJ we find:

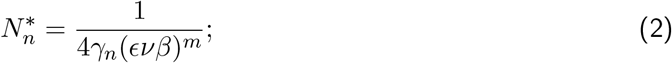

and that *γ_n_* ≈ 0.2 for all values of *m* (see Supplementary Materials for details) and we see the fit is excellent, as shown in Fig. 1. This form of *N*^∗^ indicates that once the population level rate of producing functional resistance mutants ∼ *N* (*ϵνβ*)^*m*^, which requires *m* copies of R, is sufficiently large, then the probability of resistance is large; more specifically, at the critical population size 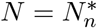, when the probability of resistance is large, the population level rate of generating functional NHEJ mutants is 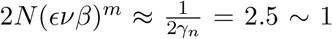 per generation. As we show in the Supplementary Information this particular form with a single fixed *γ_n_* independent of *m* arises when the rate of generation of *m*-fold resistance mutants concurrently is faster than generating them sequentially, which we show is generally the case when cleavage is efficient (1 − *ϵ* ≪ 1). To be clear, both pathways, concurrent and sequential generation of *m*-fold mutants, have a rate that scale as (*ϵνβ*)^*m*^, but they have different pre-factors determining their relative importance.

On the other hand for de novo SNPs *N*^∗^ takes the following similar functional form:

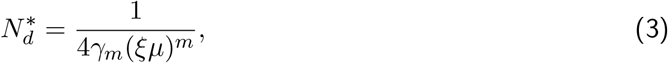

but unlike for NHEJ mutants, the fitting constant *γ_m_* is not independent of *m* and we show in the Supplementary material is of the form *γ_m_* = *s_b_τ^m^*, which arises when multi-fold resistance alleles arise sequentially rather than concurrently; here, *s_b_* is an effective/aggregate beneficial selection coefficient for the resistant haplotype with *m* functional resistant alleles and *τ* is the timescale over which these resistant mutants accumulate. By fitting the simulation data we find that *γ_m_* = {0.76, 6, 55} for *m* = {1, 2, 3}, respectively, which corresponds to *s_b_* ≈ 0.09 and *τ* ≈ 8.5 generations, which pleasingly is a time consistent with the time scale over which drive increases to high frequency. As the rate per individual of generating functional resistance at all *m* sites is (*ξµ*)^*m*^, this means the population level rate needed for the probability of resistance to be large is 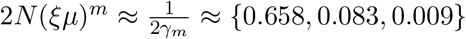 per generation, for *m* = {1, 2, 3}, respectively; this indicates that for the same individual rate of producing functional mutants at all *m* sites, the critical population size for de novo generation of mutants is smaller, and it is easier for resistance to arise de novo comnpared to NHEJ generation of mutants and the disparity increases as the number of gRNAs *m* increases.

For both NHEJ and de novo SNPs, we can summarise this information in Fig.2 and Fig.3a, by plotting contours of *p* = 0.05 on axes of *β* vs *N*, or *ξ* vs *N*, where the region to the left of the contour indicates values of these parameters for which we expect the probability of resistance to be ≤ 0.05. As indicated by Eqn.2 and Eqn.3, we see that these contours are a power law ∼ *β*^−*m*^ and ∼ *ξ*^−*m*^, respectively. The solid lines are plots of 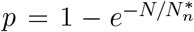 for *p* = 0.05 and *γ_n_* = 0.2 in Fig.2 and *γ_m_* = {0.76, 6, 55}, for *m* = {1, 2, 3} in Fig.3a. In both figures, the different symbols correspond to different fitness costs of mutants in the presence of wild type, and we see that here this has no effect on contours, which is intuitive as these mutants only become advantageous once drive has nearly fixed, and their effective frequency dependent selection coefficient has only a weak dependence on *σ*, the cost in the presence of wild type only.

**Figure 2:**
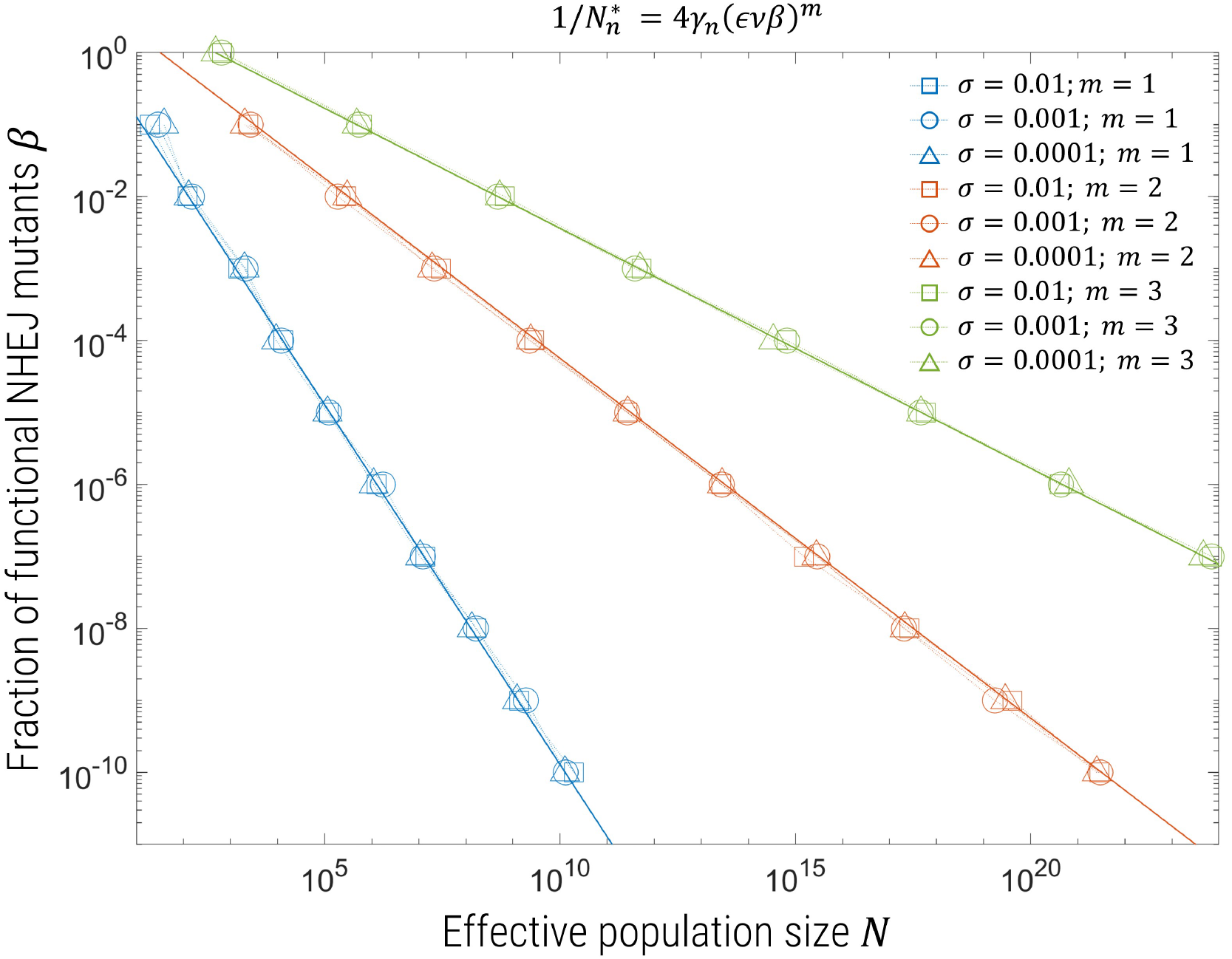
Contours for probability of resistance *p* = 0.05 as a function of *N* and *β* for *m* = 1 (blue), 2 (red) or 3 (yellow) gRNAs. Different symbols correspond to interpolated values from simulations for different fitness costs for functional mutants: square symbols *σ* = 0.01, triangle symbols *σ* = 0.001, circle symbols *σ* = 0.0001, and the solid line is a plot of 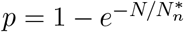 for *γ_n_* = 0.2.

**Figure 3:**
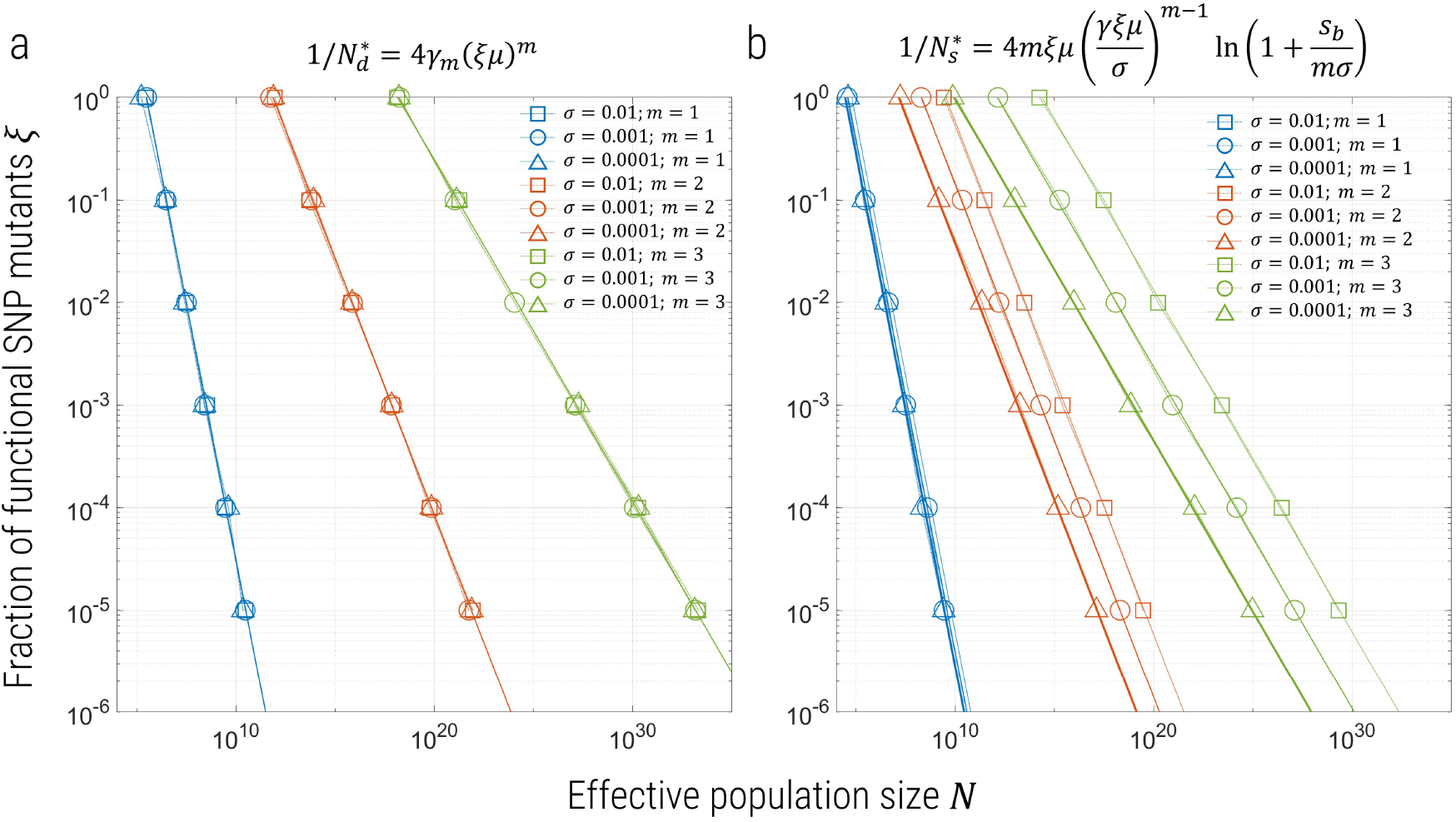
Contours for probability of resistance *p* = 0.05 as a function of *N* and *ξ* for *m* = 1 (blue), 2 (red) or 3 (yellow) gRNAs. The left panel (a) corresponds to simulations with de novo generation of SNP mutants only, while the right panel (b) are simulations with both de novo and preexisting SNP mutants. Different symbols correspond to simulations for different fitness costs for functional mutants: square symbols *σ* = 0.01, triangle symbols *σ* = 0.001, circle symbols *σ* = 0.0001, and the solid lines are a plot of *p* = 1 − *e*^−*N/N*^∗, with 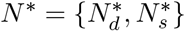 for a) & b) given by the Eqns.3 & 4, respectively, where for (a) *γ_m_* = {0.76, 6, 55} and for (b) *γ* = 2 and *s_b_* = 0.5.

### Probability of resistance: preexisting and de novo SNP mutants only

We now allow the possibility that functional resistant (R) mutants may exist in a mutation-selection balance before the introduction of drive, by running each replicate simulation for a period of time 1*/σ* before the introduction of drive, which gives sufficient time for the frequency distribution of the SNP mutants at the time of introduction of drive to have equilibrated (see Methods). As in the previous sections, we can very accurately fit curves of the probability of resistance vs *N*, using the same functional form 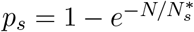, where now

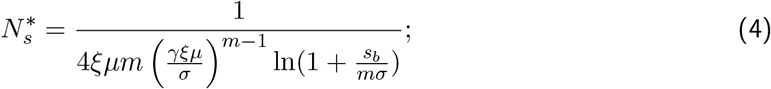

for *m* = 1 this corresponds to the theory of Pennings & Hermisson (*17*), assuming a fixed beneficial selection coefficient of *s_b_* for the resistance mutant in the presence of the drive allele, which is derived by calculating the average probability of fixation over the distribution of allele frequencies in mutation selection balance, where the fitness cost is *σ*, before drive is introduced. For sufficiently strong selection 4*Nσ* > 1, this allele frequency distribution is closely approximated by a gamma distribution with shape parameter *θ* = 4*Nξµ* and rate parameter *α* = 4*Nσ*. In the case of *m* > 1 gRNAs, the functional form for *N*^∗^ can be heuristically motivated by assuming the frequency distribution is of the same form (gamma distributed), but with a modified rate parameter *α_m_* = 4*Nmσ* and shape parameter 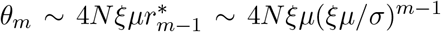, where 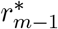 is the frequency of mutants with *m* − 1 R alleles and a single W in mutation-selection balance before drive is introduced. For example, if there are *m* = 2 gRNAs then 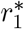 is the frequency of RW, while for *m* = 3 RNAs *r*_2_ is the frequency of RRW. We show in the Supplementary material that 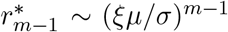. *θ_m_* represents the effective mutational flow from *m* − 1 mutants to *m* mutants, which is balanced by negative selection, where each *m* mutant has fitness cost *mσ*, before the introduction of drive. By averaging the probability of fixation over this distribution we recover the scaling law for 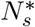 stated above. We do not attempt to calculate an exact theory, but use a fitting parameter *γ* to represent the scaling between these heuristic considerations and an exact theory.

We first fit the curves of probability of resistance vs *N* for *m* = 1, which gives an approximate value of *s_b_* ≈ 0.5 for all values of *ξ* (Supplementary Material), where *γ* does not appear in 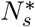. For *m* > 1, we fix *s_b_* to this value and fit the curves of *p*(*N*) for the single fitting parameter *γ*, and find that *γ* ≈ 2 for both *m* = 2 and *m* = 3, as shown in the Supplementary material. In all cases, we find the curves fit the simulation data very well.

We summarise these results by plotting contours of constant probability of resistance *p* = 0.05 as a function of *ξ* and *N* as shown in Fig.3b; regions to the left of each curve represent combinations of *ξ* and *N*, for which the probability of resistance is less than 5%. The major effect of preexisting mutations is to very greatly reduce the effective population size needed before resistance arises for more than *m* > 1 gRNAs. Importantly, unlike for de novo SNPs or NHEJ mutations, there is a significant effect for changing selection, where the probability of resistance increases for decreasing *σ*, because less harmful mutations will segregate at a higher frequency before release. Though Hermisson and Pennings (*17*) showed this to be the case for a single site (*m* = 1) (*6, 17*), the effect, as we see is only weak and logarithmic in *σ*; here however, for *m* > 1, *N*^∗^ ∼ *σ*^*m*−1^, which represents a significant amplification of more weakly selected standing variation.

### Probability of resistance: NHEJ, de novo and preexisting SNPs combined

Finally, we combine all three mechanisms by which resistance can arise to assess their relative importance as a function of varying *β*, *ξ* and *σ*. The summary of these results are shown in Fig.4, for contours of *p* = 0.05 for *β* vs *N*, for the case of a) all SNPs being functional (*ξ* = 1) and b) 1% functional SNPs (*ξ* = 0.01). The same broad trends as observed in the simulations of the previous sections are seen, where decreasing *β* increases the range of population sizes for which resistance is improbable. However, in the presence of preexisting SNPs, we find that for sufficiently small *β*, this effect plateaus and the 5% contours are constant with respect to *N*, as the dominant and more rapid mechanism of resistance becomes SNPs from standing variation. We also find that increasing *ξ* decreases the population size at which resistance arises at 5%, given by a uniform shift of these curves to the left, as seen by comparing panel a) to b). The solid lines in each plot correspond to assuming the probability of resistance from NHEJ mutations and preexisting (& de novo) mutations are independent: *p_res_* = 1 − (1 − *p_n_*)(1 − *p_s_*) = 1 − *e*^−*N/N*∗^, where

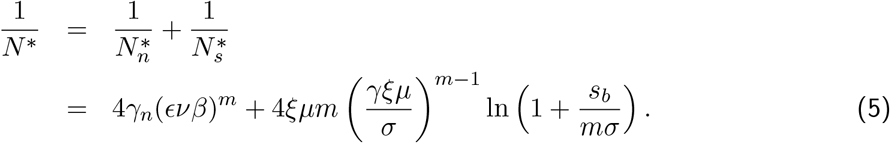

**Figure 4:**
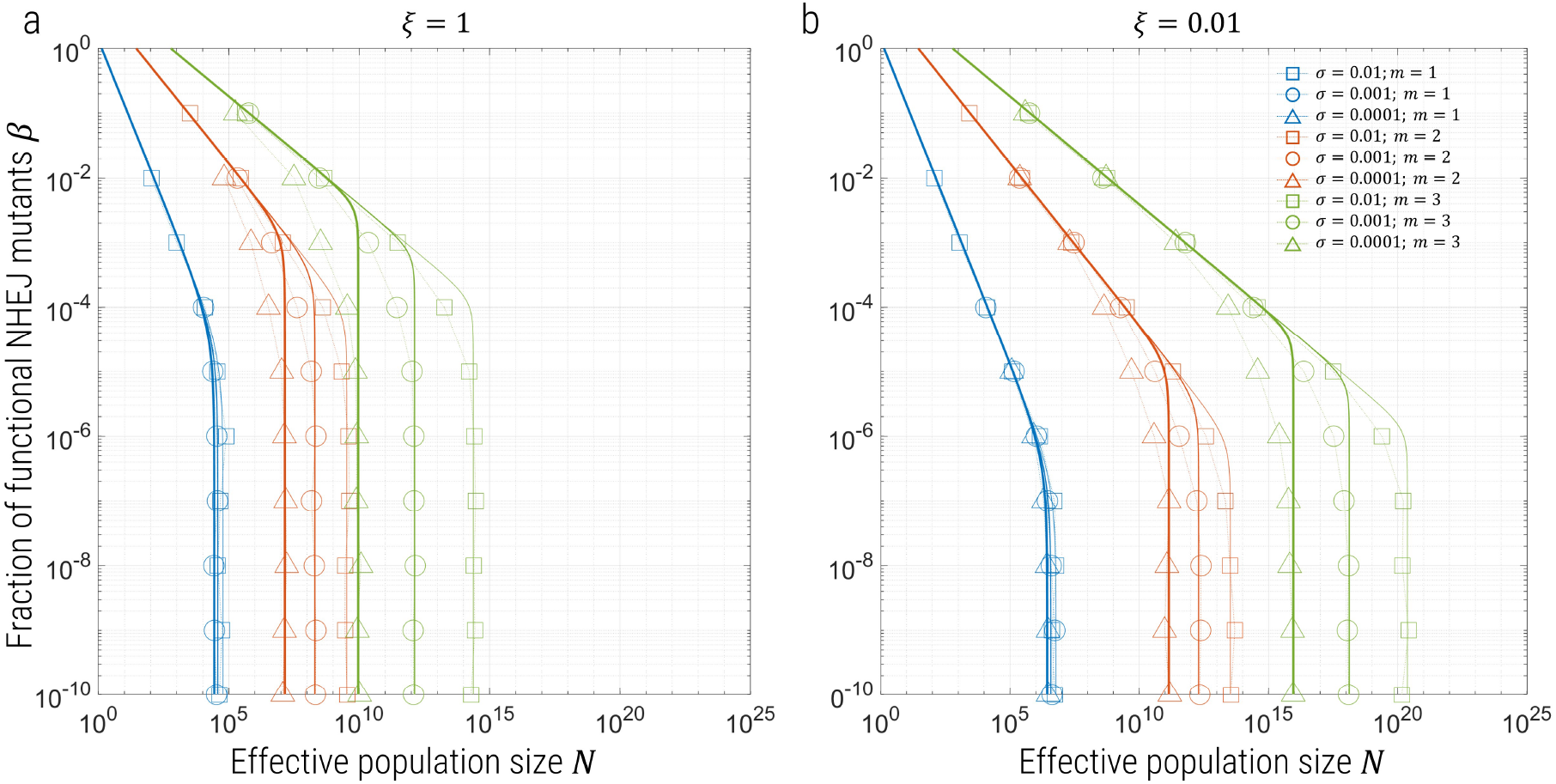
Contours for probability of resistance *p* = 0.05 for *β* vs *N* and a) *ξ* = 1 and b) *ξ* = 0.01, for simulations including NHEJ, de novo and pre-existing SNPs. Blue symbols represent *m* = 1, red *m* = 2 and yellow *m* = 3, where the different symbols present different fitness cost *σ* of mutants in the presence of wild type, as shown in the legend. The solid lines are plots of 1 − *e*^−*N/N*∗^, for 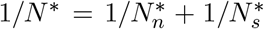 — which assumes the probability of resistance from NHEJ and SNPs are independent — for exactly the same values of the *γ* parameters in the previous sections/plots, where the thick lines correspond to *σ* = 0.0001, intermediate thickness to *σ* = 0.001 and thin lines to *σ* = 0.01.

The plots use the parameters *γ_n_* = 0.2 and *γ_s_* = 2, as derived from fitting in the previous sections, where the different line thicknesses correspond to different values of *σ*; we see that asymptotically for either 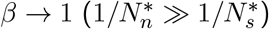 or 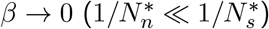, where NHEJ mutations are dominant and SNPs are dominant respectively, the solid lines match the simulation data very well, as expected. However, for the intermediate regime where NHEJ mutations and SNPs from standing variation are equally dominant, there is a mismatch, which indicates the assumption that resistance from NHEJ and SNPs are independent is relatively poor. This is as expected, since this heuristic ignores the mosaic nature in which resistance arises at multiple target sites, where for example at one site resistance may have arisen by standing variation and the other two by NHEJ; nonetheless we see this heuristic gives a good guide to the probability of resistance in the presence of both NHEJ and standing SNP variation.

### How many gRNAs are needed to prevent resistance?

An important practical question that arises is how many gRNAs is sufficient to prevent resistance to a high probability. Assuming a regime of *β*, where NHEJ mutations are dominant, using Eqn.1 & 2 this result we can find the required number of gRNAs to keep the probability of resistance to less than or equal to *p*:

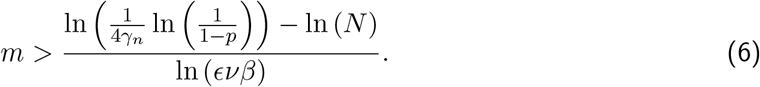

This result is similar to Equation 2 of (*7*), the main difference being that it is derived by approximately solving the dynamics of resistance allele generation, using their probability of fixation (Supplementary Information) and explicitly accounting for a fraction *β* of functional NHEJ mutants.

We can find an equivalent expression for de novo SNPs using Eqn.3, but since our results indicate that standing variation will always be at least as important as de novo generation of SNPs, we directly consider the constraint on *m* for standing variation. Using Eqn.1&4, we can in principle calculate the minimum number of gRNAs *m* required to prevent resistance to a specified probability. Eqn.4 is transcendental, but if we replace the weak *m* dependence in the logarithm 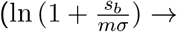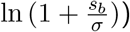 then for *σ* ≫ *ξµ*, we find the approximate expression:

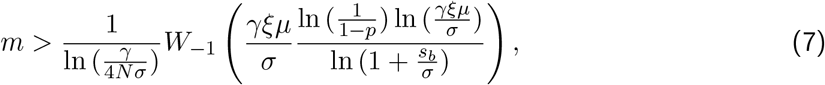

where *W_k_*(*x*) is the Lambert *W* function, which are solutions of the equation *we^w^* = *x*.

When both mechanisms of resistance are possible, the effective critical population size *N*^∗^ is a combination of 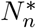 and 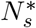, as shown in Fig.4 Eqn.5 is a reasonable approximation, but it is difficult to invert this expression to find how many gRNAs are needed to prevent resistance to a certain probability *p*. Here, we instead estimate this critical population size for different values of *m*, *β*, *ξ* and *σ* by reading off Fig.4 or by extrapolation using the approximation in Eqn.5, which we can compare to whichever target population size we have in mind for an application of drive. We can examine two different extremes: 1) *β* = 10^−4^, which corresponds to assuming functional NHEJ mutants are quite rare, as we might expect given the expectation that NHEJ will tend to produce significant genetic changes like multiple base-pair insertions and deletions and 2) *β* = 10^−2^, which is more pessimistic, should in fact NHEJ more readily produce less significant genetic changes, which are more likely to be functional. For the fraction of functional SNPs, we assume a worst case that *ξ* = 1 and the scenario that *ξ* = 0.01, which roughly corresponds to on average a single functional mutant in all the 3*L* one-step mutants about the wild type sequence in a target site of size *L* = 18bp (values of *ξ* ≪ 0.01 effectively correspond to *ξ* = 0). To calculate *N* corresponding to when resistance is equal to 5%, we can read off from Fig.4 or use our extrapolation of the simulations using Eqn.5 to different values of *σ*. These population sizes are shown in Table.1.

**Table 1:**
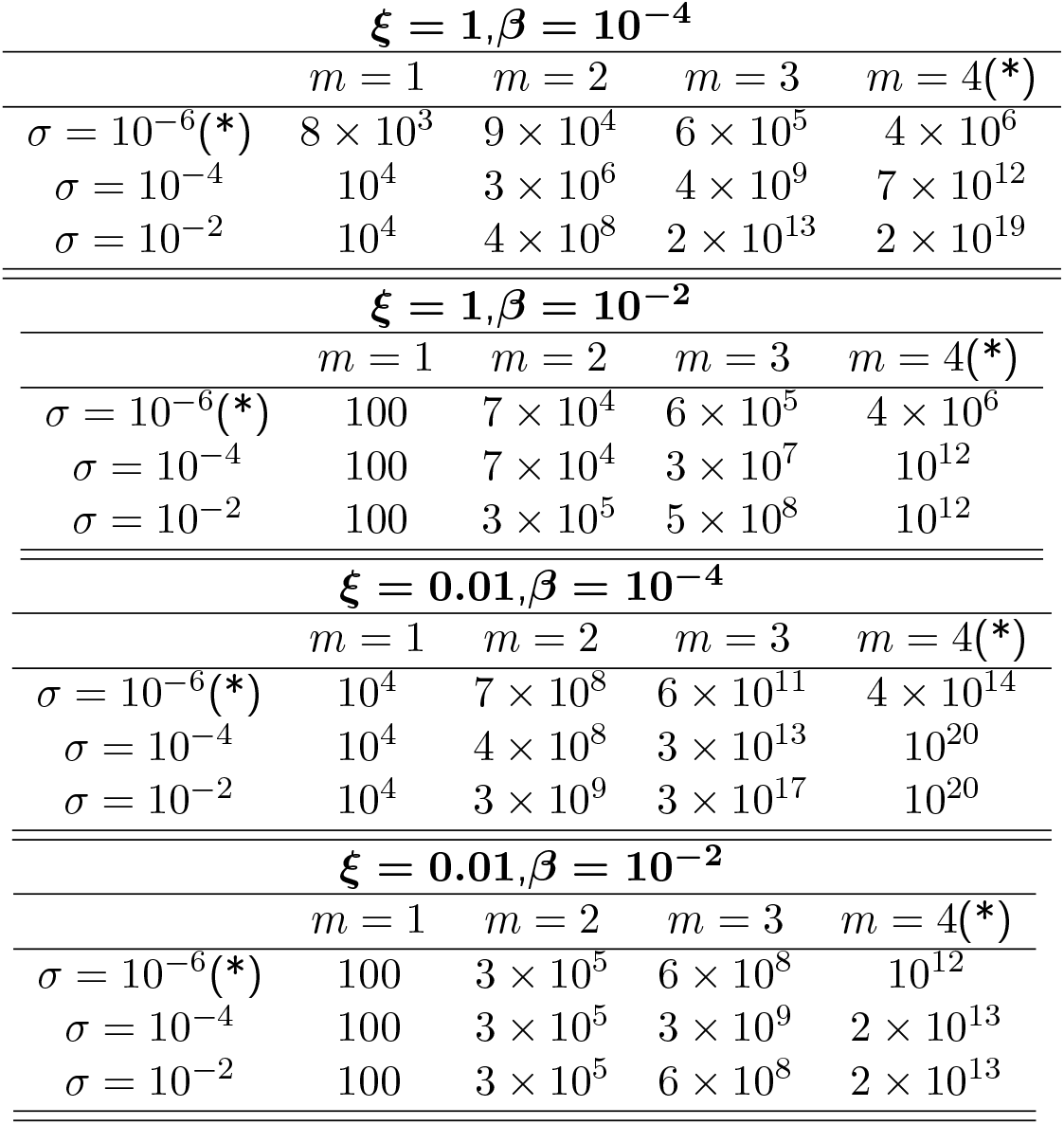
Effective population sizes *N* at which a probability of resistance *p* = 0.05 is obtained from simulations and theory for different numbers of gRNAs *m* and different selection coefficients *σ*, for the fixed value of fraction of functional mutants *ξ* = 1 and *ξ* = 0.01. The starred values indicate population sizes extrapolated from simulations results using heuristic theory (Eqn.5).

### Time to Resistance

In the Supplementary material we also examine the mean time taken for resistance to arise, *given* that resistance arose, where resistance corresponds to the establishment and then fixation of a functionally resistant allele. We find across all simulations, very consistently, the mean time is between 70 and 100 generations, where as expected in general for large values of *β* we have shorter times, since the rate of producing NHEJ mutants is larger and as the population size is increased the time to resistance increases. Broadly, we find that if variants are more deleterious before the introduction of drive and/or as the population size increases, that the time to resistance is longer. These results point to the fact that the time for resistance to arise is conditioned on resistance arising, and so all the variants that are destined to give resistance, must be generated, or pre-exist, before the population is eliminated; population elimination happens on a timescale set by the dynamics of drive replacing the wild type — which does not change for different simulations performed in this paper — and by the size of the initial population, where larger populations take a logarithmically longer time to elimination (on average).

## Discussion

Understanding and overcoming resistance in suppression drive systems is a major obstacle to successful control of natural populations that are vectors for disease such as malaria (*3, 4*). As this work and previous research shows, population size is a key determinant of the probability of resistance (*5–7*). In particular, we have highlighted the role of the critical population size *N*^∗^, as a useful summary measure of the probability of resistance compared to some target natural population size. Multiplexing of guide RNAs, such that resistance is required in all gRNAs, is a promising anti-resistance strategy, as it aims reduce the individual rate of resistance sufficiently that *N*^∗^ is much greater than any realistic population size. Our results have highlighted 5 key parameters determining *N*^∗^ and the probability of resistance evolving for multiplexed drive: *β*, *ξ*, *σ*, *m*, and *N*. The expressions we derive from simulations for *N*^∗^ in terms of these parameters inform on the relative importance of non-homologous end-joining vs de novo single nucelotide mutations vs standing variation of SNPs. Our key novel finding is a significant amplification of the role that weakly deleterious standing genetic variation plays in determining resistance in multi-site evolutionary systems, compared to de novo mutation, such that resistance is probable even when the number of gRNAs is large and for not very large population sizes. In other words, the intuition that many gRNAs are guaranteed to prevent resistance can potentially fail if standing variation is not taken into account.

Researchers developing gene drive constructs for population suppression will have some level of control over all of the key parameters identified above, but the last of these, by choice of target site. NHEJ typically produces insertion or deletion mutations (*3*), so choosing a target site that is unable to tolerate length variation will be one way to reduce *β*, and this would usually be a top priority in choosing a target site. Hammond et al. (*3*) have demonstrated that having a single target site that can tolerate length variation quickly leads to the evolution of resistance, even in small populations (*N* ≈ 600). Though data are scarce, the next most frequent type of mutation produced by NHEJ is presumably single nucleotide changes at the cut site, and ensuring those are nonfunctional would be a second priority. Assuming a target population of *N* = 10^6^ and the baseline parameter values, Eqn.6 indicates that under the worst case scenario of *β* = 1, *m* = 6 gRNAs will be needed to have the probability of resistance arising due solely to NHEJ be less than 0.05, whereas if *β* can be reduced to 0.01, then only *m* = 3 will be needed, and if *β* can be reduced further to 10^−4^, then only 2 will be needed. If the target population is *N* = 10^9^, then the corresponding values are *m* = {8, 4, 2} gRNAs for *β* = {1, 0.01, 10^−4^}, respectively.

Single nucleotide mutations can also arise spontaneously, and, if the population is large, may be already present before release. These may also provide resistance against cleavage, particularly if they are near the cut site (*19*). The likelihood that resistance evolves from these mutations depends on *ξ*, the fraction of them that are functional (since it is only these that can spread through a population) and on *σ*, the extent to which those functional resistant mutations reduce fitness (since that affects their frequency in the pre-release population). Again, all else being equal, more functionally constrained sequences will be preferred. In principle, genetic surveys of sequence variation at the target site may provide useful information on *ξ* and *σ* and the probability of resistance evolving through standing variation. Assuming a target population of *N* = 10^6^, baseline parameter values and *σ* = 10^−6^, then using Eqn.7 under the worst case scenario of *ξ* = 1, *m* = 4 gRNAs are needed to ensure the probability of resistance evolving from de novo mutations and pre-existing variation is less than 0.05, whereas just *m* = 2 gRNAs are needed if *σ* = 10^−4^ or *σ* = 0.01, which demonstrates the sensitivity to weakly deleterious standing variation. However, if *ξ* = 0.01 then only 1 gRNA is needed, irrespective of the value of *σ*. If the target population is *N* = 10^9^ the corresponding values are *m* = {7, 3, 2} gRNAs are needed respectively, for *ξ* = 1 and *σ* = {10^−6^, 10^−4^, 0.01}, whereas for *ξ* = 0.01, *m* = 2 gRNAs are needed, regardless of *σ*.

An important application of suppression drives is to control populations of mosquitoes to reduce the burden of malaria on human populations. Recently, the contemporary effective population size for *Anopheles gambiae* in sub-Saharan Africa was estimated as *N* ∼ 10^9^, using a new method based on analysing soft sweeps (*9*). The above considerations of NHEJ and standing variation separately give an indication of how many gRNAs are required to achieve a probability of resistance less than 0.05, in each scenario, but including both mechanisms, we refer to Table.1. Given a target population size of *N* = 10^9^, and assuming *ξ* = 1, these numbers indicate that if resistance alleles are strongly deleterious (*σ* = 0.01) to moderately deleterious (*σ* = 10^−4^) then resistance can be prevented with *m* = 3 gRNAs if *β* = 10^−4^, or *m* = 4 gRNAs if *β* = 10^−2^. However, if the resistance alleles are very weakly deleterious (*σ* = 10^−6^) then even 4 gRNAs is not sufficient, for both values of *β*, which exemplifies the strong amplification of the probability of resistance in the presence of weakly selected standing variation. On the other hand, if *ξ* = 0.01, which roughly corresponds to on average a single functional mutant in all the 3*L* one-step mutants about the wild type sequence in a target site of size *L* = 18bp, we can prevent resistance for all values of *σ* and *β* with *m* = 3 gRNAs, except for very weakly deleterious mutants (*σ* = 10^−6^) and *β* = 10^−2^, which requires at least *m* = 3 gRNAs. These considerations highlight the need to empirically determine both *β* and *ξ* for putative target sites for multiplex suppression drive applications, using genetic screens which determine the functionality of NHEJ mutants and by examining SNP variants and their fitness effects from population genomic data.

These simulations make a number of simplifying assumptions that are necessitated by modelling the already relatively complex situation of multiplex drive. We assume that all of the nucleotides *ξ* corresponding to the fraction of functional mutations in each target site, once mutated completely abolish cleavage and give complete resistance. These assumptions can be relaxed in various ways, such that each mutation contributes to partial reduction in cleavage efficiency; within our model this would correspond to a different effective value of *ξ*. We also assume there is no recombination between sites and related to this that resistance only arises due to target site effects; recombination and off-target resistance has been explored using deterministic modelling (*20, 21*). We also have not explored the role of varying a number of parameters like cleavage efficiency *ϵ*, the intrinsic growth rate *R_m_*, or the fitness parameters of drive; some of these have been previously explored in the context of single gRNAs (*6, 7*) and these works and ours in general indicate these will have a secondary role compared to the population scaled rate of NHEJ and de novo mutation. For example, increasing *ϵ* will tend to increase the rate that drive replaces wild type, and we may expect one consequence is a quantitative reduction in the probability of resistance as there is less time to generate mutants and establish before population elimination, but the qualitative results we have presented will not change. However, there is evidence that the overall effective efficiency of suppression may diminish when there are large numbers of gRNAs (*22*). There is also the possibility that homing events may be more error-prone than normal DNA replication and could lead to the loss of function of one or more gRNAs, which we leave to future modelling efforts, but could increase the probability of resistance evolving depending on the rate of such events occurring.

On one hand, it is not clear that spatial structure will strongly affect these results, since the population size we consider here should be very close to the census size; whether spatially separated or in a well-mixed system, the number of mutations arising per generation will depend on the total number of individuals in the population. However, what is likely to be different is how quickly a resistance mutation establishes and spreads; in spatially structured populations selection is effectively weaker (*23*), reduced by factor 1 − *F_ST_*, where *F_ST_* = 1*/*(1 + 4*Nm*) is Wright’s fixation index for the island model, which means fixation would be less rapid in a geographically dispersed population. This could lead to an increase in the critical population size *N*^∗^, since at a given population size the probability of resistance is smaller, as elimination is more likely before the resistance allele becomes sufficiently prevalent in the population. However, this is likely to only have significant effect in very highly structured populations (very limited migration), since typically selection for resistance mutants is very large once drive has risen to large frequency; however, if spatial structure is important, it would mean a systematic underestimation of the critical population size *N*^∗^. Importantly, in models with spatial structure even arbitrarily strong gene drives may not eliminate a target population (*24–26*). This effect can arise if the gene drive causes reductions in population density, which leads to increased inbreeding, which in turn reduces the efficacy of the drive (*27,28*). If the population is not eliminated, then eventually one would expect resistance to evolve, though if the population is substantially suppressed this may take a long time.

These results also have broader implications for evolutionary theory, particularly evolutionary mechanisms by which adaptation occur in response to an environmental change, such as the introduction of drive, or in other contexts, such as resistance to antibiotics or vaccines. Theory for a single site (*m* = 1) in various evolutionary contexts includes the question of which is more important, de novo vs standing variation for adaptive evolution (*17, 29*), population rescue (*14*) or in the context of the evolution of gene drive resistance (*6*). All these studies show that changing the magnitude of the fitness cost *σ*, before the environmental change, has a relatively weak effect on the probability of resistance, as borne out by the logarithmic dependence of 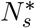 on *σ* for *m* = 1 in Eqn.4. However, a major new finding is that for the multiplex drive case, where resistance alleles must evolve at all *m* target sites in order for resistance to arise, there is in fact a marked dependence on the fitness of resistance alleles before the introduction of drive. This arises from a complex mutation-selection-drift balance between fully resistant (*m* R alleles) and incomplete resistant (less than *m* R alleles) haplotypes, resulting in a significant amplification of weakly deleterious alleles in their contribution to resistance in a multiplex scenario, and a significant reduction in the critical population size with standing variation compared to de novo mutation, as seen in Fig.3b. We can quantify this amplification by calculating the ratio

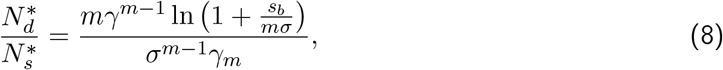

which has values 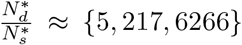 for *m* = {1, 2, 3} and *σ* = 0.01, and 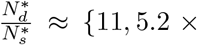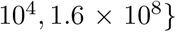 for *σ* = 10^−4^, which are very large amplification factors for *m* > 1 and with consequent implications for the prediction of the probability of resistance for multiplexed gene drives (Note that these very large differences are somewhat hidden in Fig.3 as the results are plotted on a log scale and over many orders of magnitude).

This finding and Eqn.4 should have general applicability to evolution of multi-site resistance in many different contexts (*30*). This includes multi-drug resistance to antibiotics (*31, 32*), anti-viral therapies (*33*) and cancer (*18, 34*). The theory is particular applicable to the evolution of antibody/vaccine escape mutants, where multiple epitopes on a virus structure are targeted by antibodies to reduce infection, and where recombination is likely to play a small role. Although each of these evolutionary scenarios carry with it particular idiosyncracies, which would require proper attention, qualitatively, it is clear that very weakly deleterious standing variation can very easily give rise to small critical population sizes even for moderately large *m*.

A key case in point is the evolution of vaccine/antibody escape of Sars-Cov2, and viruses in general, where a large number of epitope sites are targeted by neutralising antibodies (*35–40*). If infections are persistent, as currently with Sars-Cov2 and other endemic diseases, we do not strictly have a population rescue problem, but Eqn.4 can be viewed as the critical population if a long time was allowed for standing variation to develop. In the case of Sars-Cov2 as many as *m* = 20 (*38*) epitope sites may need to evolve to prevent antibody binding, whilst for measles only *m* = 5 may be required (*40*). The results here (Eqn.4) would broadly suggest that higher levels of infection could increase levels of standing variation and make vaccine escape very probable. Although it is too early to assign specific selective causes, if it is the escape of natural or vaccine-induced immunity that has led to the recent emergence of the B.1.1.529 (Omicron) variant with roughly 30 mutations in the spike-ACE2 receptor binding protein, then these mutations could potentially have accumulated via this mechanism, prior to whatever change in selective pressures that allowed its establishment; this may have occurred in a single prolonged infection in an immunocompromised individual, and/or through chains of transmission between a number of acutely infected individuals. We can understand qualitatively how such a large number of mutations *could* arise, if they were weakly deleterious, even for moderate effective populations sizes, since we can re-arrange Eqn.4 (ignoring the weak logarithmic term and assuming 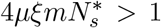) to give 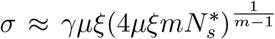, which for large *m* decreases extremely slowly; in other words the value of *σ* required for say *m* = 30 mutations to give rise to resistance will be marginally smaller than that required for *m* = 10. However, the applicability to Sars-Cov2 is arguable as the time since the start of the pandemic may not be sufficient for multi-site variants to reach their equilibrium frequency, although this modelling and theory provides a framework to address this question.

Overall, our results provide a foundation to understand how resistance arises in multiplexed suppression-drive systems and the paramount role that standing variation plays in greatly amplifying weakly deleterious variation to give resistance at even moderate population sizes for a large number *m* of gRNAs. The results highlight the need to characterise important unknown parameters such as the fraction of functional mutants at drive target sites, due both to non-homologous end joining mutations and single nucleotide mutations, which can significantly affect the probability that resistance arises. Finally, we suggest the multi-site evolutionary rescue problem studied here and the role of standing variation has general applicability to multi-drug antibiotic, anti-viral and cancer resistance, as well as the evolution of vaccine escape.

## Methods

We use non-spatial Wright-Fisher stochastic simulations of drive with separate sexes throughout, but with coupling to population dynamics using density-dependent Beverton-Holt growth. The details of these simulations are given in the Supplementary Information. The simulations are stochastic, even when population sizes are very large, because resistant mutations always arise at small frequency, so genetic drift needs to be explicitly considered. Resistance corresponds to the establishment and then fixation of functionally resistant alleles, which we define to be a frequency greater than 0.95. The simulations entail stochastic dynamics with one, two, or three gRNAs, where at each *target* site there are 4 alleles that can occur W (wild type), D (drive), R (functional resistance), N (non-functional resistance); this means that across an *m*-fold target site, we have an *m*-locus, 4-allele population genetic system with no recombination. However, as we assume the drive construct is copied over as a whole, in practice, it is an *m*-locus 3-allele system +1 for the D allele. On a single chromosome the possible haplotypes (assuming no positional effects) for a 2-fold system are WW, WR, RR, WN, RN, NN, DD, which is a total of *n*(*n* + 1)*/*2 + 1 = (3 × 4)*/*2 + 1 = 7 haplotypes, and an analogous calculation in the Supplementary Information for *m* = 3 gives a total of 11 haplotypes.

The alleles are assumed to have no effect on male fitness, while fitness effects in females are as follows. W alleles have zero fitness cost, and D and N alleles are deleterious with homozygous fitness cost *s* = 1 and heterozygous fitness costs (when paired with a W allele) of *h* = 0.3 and *h_N_* = 0.02, respectively; the larger dominance coefficient for drive represents potential somatic fitness costs to heterozygotes females, from leaky expression of the Cas-9 protein (*4*). The heterozygous fitness cost of R, before the introduction of drive, is *σ*, which we vary in the simulations. When combined in haplotypes we assume each target site has independent fitness effects, which means a single occurrence of an N allele is deleterious. Fitness costs are manifest as reduced female survival. Further details are given in the Supplementary Information.

In W/D heterozygotes (and their multiplexed equivalents discussed in the Supplementary Information) we assume a cleavage efficiency *ϵ* = 0.95, and a non-homologous end joining (NHEJ) rate of *ν* = 0.05 approximately representative of the target site in the gene *doublesex* in *Anopheles gambiae* (*4*). We assume NHEJ mutants produce functional resistant alleles R with probability *β* and nonfunctional resistant alleles N with probability 1 − *β*. As a result D gametes are generated from conversion of W at a rate (1 − *ν*)*ϵ*, functional resistance alleles R at rate *ϵβν*, and non-functional alleles N at rate (1 − *β*)*ν*, while the fraction that remain wild type is 1 − *ϵ*.

We assume the probability of cleavage at each available target site given by efficiency *ϵ*, occurs independently at different sites, so that the fraction of non-driving gametes produced from a genotype with a non-driving allele/haplotype paired with a driver is 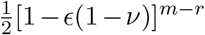, where *m* is the number of gRNAs and *r* ≤ *m* is the number of resistant (R or N) sites in the non-driving allele. Thus when there are multiple gRNAs the presence of a resistant sites give some protection to the chromosome from being cut, even if one or more cleavable sites remain.

In addition, functional resistance alleles R are generated de novo at rate *ξµ*, where *µ* is the mutation rate for the length of site of interest and *ξ* is the fraction of SNP mutations that are functional, and non-functional resistance alleles at rate (1−*ξ*)*µ*. We assume *µ* = 18*µ*_0_ = 5.4×10^−8^, where 18 is the length of each target site and *µ*_0_ = 3 × 10^−9^ is the base-pair mutation rate measured for *Drosophila* (*41*).

### Standing genetic variation

Before drive is introduced into a population there may be pre-existing functionally resistant SNPs in the population which would affect the probability of resistance arising. To study the effect of standing variation we run replicate simulations where we allow a burn-in period of 1*/σ* generations to allow for the population to come to a mutation-selection balance equilibrium. The initial frequency of the various resistance alleles/haplotypes when drive is introduced at *t* = 0 is then implicitly drawn from the mutation-selection balance equilibrium. Note that in the case of *m* > 1 the mutation-selection balance distribution will be complex with different frequencies for haplotypes carrying different numbers of resistance mutations R.

### Gaussian-Poisson hybrid approximation to generate multinomial random numbers

In this paper we run simulations to very large effective population sizes. Whilst it is typical in such a scenario to ignore the stochastic part of the evolutionary dynamics by using deterministic dynamics, this is only accurate if the allele frequencies are large themselves, or equivalently the number of copies in the population are large (≫ 1). We are interested in the dynamics of resistance, which by definition means we need to study situations where the allele arises by de novo mutation as a single copy in a single individual, where it must survive genetic drift, or exist at very low frequency as standing variation. When there are multiple alleles, particularly when simulating Wright-Fisher evolutionary dynamics, this is accomplished simply by drawing multinomial random numbers. However, when the effective population size is large this can become increasingly slow. In addition, the maximum population size is restricted to the largest integer that can be stored in a computer; for the GNU scientific library’s implementation of multinomial random number generators, this is limited to 32-bit, which gives a limit of roughly 4 × 10^9^.

An alternative approach is to use the multivariate Gaussian approximation to the multinomial distribution:

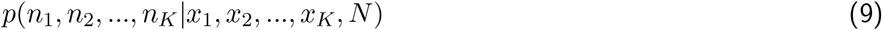

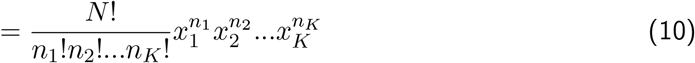

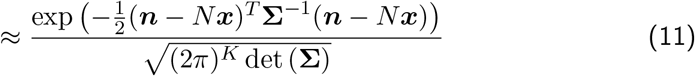

where ***n*** = (*n*_1_, *n*_2_, …, *n_K_*)^*T*^, ***x*** = (*x*_1_, *x*_2_, …, *x_K_*)^*T*^, are the vectors of the numbers drawn of *K* alleles, and their expected frequency, respectively, and **Σ** is the scaled covariance matrix of the multinomial distribution, where 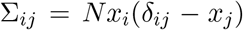. However, this approximation is poor when for any of the alleles 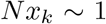. In this limit, these rare alleles are well-approximated by a Poisson distribution. The approach taken in this paper is therefore to partition the alleles into a rare category 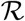, if 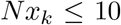, and non-rare if 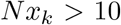, where the former is drawn from independent Poisson distributions, while the latter from a multivariate Gaussian distribution conditioned on a smaller total population size 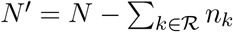:

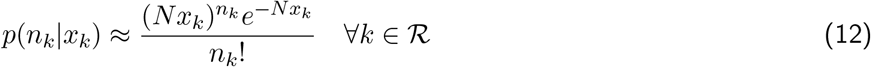

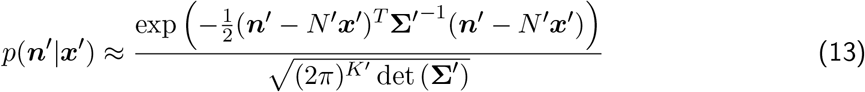

where the vectors ***n***^′^ and ***x***^′^ only take elements 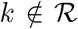, whose length is 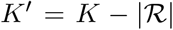, and the covariance matrix 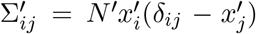. We can assume independent Poisson distributions for each of the rare alleles precisely because they are rare and the effects of drift are approximately independent of each other, and the constraint of constant population size is imposed on the non-rare alleles through the modified population size *N* ^′^ and the correlation structure in the covariance matrix **Σ**^′^.

## Acknowledgments

We acknowledge useful discussions with Ioanna Morianou on the genetics of resistance and comments on the manuscript by John Connolly and John Mumford. This work was supported by grants from the Bill & Melinda Gates Foundation and the Open Philanthropy Project.

## Supplemental Materials

### Methods

#### Single guide RNA

Simulations with a single guide RNA (gRNA) or target site require specification of a 4 × 4 fitness matrix for both males ***W***_m_ and females ***W***_f_, whose elements *w*_*ij*_ are the fitness of the genotype *i/j*, where for example genotype 1/3 ≡ W/N. In the absence of mutation and drive the allele frequency vector in next generation of females ***x***_*t*+1_ and males ***y***_*t*+1_ can be expressed succinctly in matrix notion in terms of the frequency vectors in the current generation ***x***_*t*_ and ***y***_*t*_ as follows

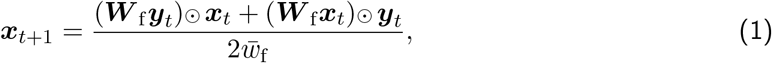

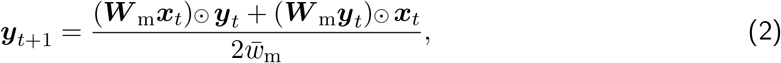

where the mean fitness of males and females is given by the quadratic forms

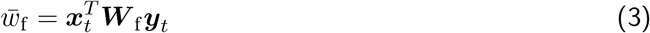

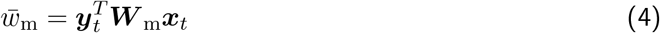

and where the ⊙ operation is the element by element (Hadamard) multiplication operation, i.e. ***z*** = ***x*** ⊙ ***y*** is equivalent to *z*_*i*_ = *x*_*i*_*y*_*i*_.

The fitness matrix for males, ***W***_m_ is a 4 × 4 matrix, where each entry, (*W*_m_)_*ij*_ = 1. For females, we assume all functional genotypes (exc N and D) have fitness 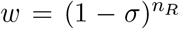, where *n*_*R*_ is the number of R alleles in the genotype, so *w*_11_ = *w*(W/W) = 1, *w*_12_ = *w*_21_ = *w*(W/R) = (1 – *σ*)

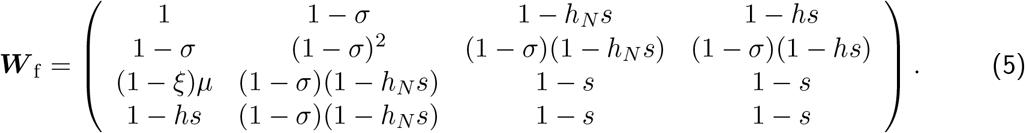

and *w*_22_ = *w*(R/R) = (1 – *σ*)^2^. On the other hand for non-functional heterozygotes, we assume a dominance coefficient *h* with drive to represent somatic leaky expression affecting viability, with a typical value *h* ≍ 0.3, and a very recessive dominance coefficient (*h* _*N*_ = 0.02) with non-functional resistance alleles. W/D heterozygotes; heuristically, this leads to the fitness of genotypes that include functional and non-functional alleles of the form 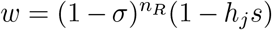, where as before *n*_*R*_ is the number of R alleles in the genotype and if *j* = N, *h*_*j*_ = *h*_*N*_ and if *j* = D, *h*_*j*_ = *h*. Genotypes with only non-functional alleles N and D are very deleterious and have fitness *w* = 1 – *s*, where *s* = 1. Altogether, this gives the following fitness matrix ***W***_f_ for females:

Including mutations is straightforward using a mutation probability matrix ***M***, whose elements *M*_*ij*_ represent the probability of generating allele *i* per generation from allele *j*, where this stochastic matrix, which acts on probability/frequency *column* vectors must have the property Σ_*i*_ *M*_*ij*_ = 1:

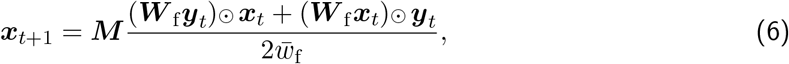

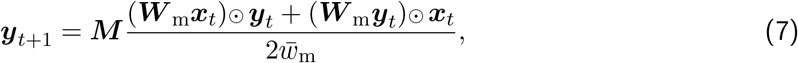

We make the assumption that there are only two mutational paths W → R and W → N, which happen with rate *ξμ* and (1 – *ξ*)*μ*, respectively and there are no backmutations to W (Fig.1. This gives a mutation matrix:

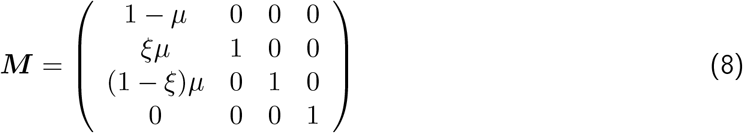

Finally, we add the effect of drive by following (1), where we assume drive acts during meiosis after viability selection. Drive is only produced by heterozygotes W/D, whose frequency after selection and mutation is 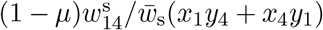 for females and males, where s = {f, m}, respectively, which gives the following allele frequency update equations

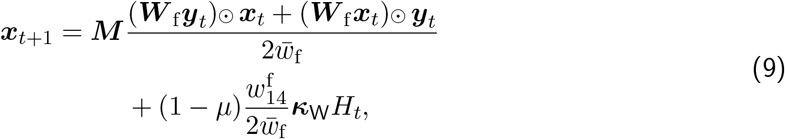

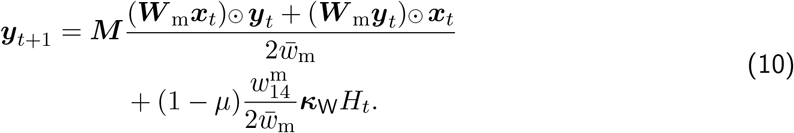

where *H*_*t*_ = (*x*_1_)_*t*_(*y*_4_)_*t*_ + (*x*_4_)_*t*_(*y*_1_)_*t*_ is the drive-heterozygote frequency (before viability selection acts) and the vector ***κ***is stoichiometry like vector describing the change in fractions of each alleles produced as gametes from the W/D heterozygote relative to no drive:

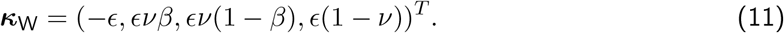

Eqns 9 & 10 are then treated as the mean or expected frequency in a Wright-Fisher multinomial sampling process. To allow for fluctuations in relative numbers o f m ales and females, we perform Wright-Fisher sampling with total population size constraint *N*_*t*+1_ on the population vector of female and male allele frequencies 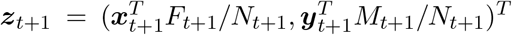, where now ***z***_*t*+1_ is a 8 column vector, and for example *z*_3_ is the frequency of female R alleles, and *z*_5_ is the frequency of male W alleles, where *F*_*t*+1_, *M*_*t*+1_ and *N*_*t*+1_ are the total number of females, males and their sum, which are calculated using a Beverton-Holt scheme described next.

To couple the population genetics calculated here to the population dynamics, we use the Beverton-Holt model of density-dependent growth for male and female populations separately, assuming only females give birth:

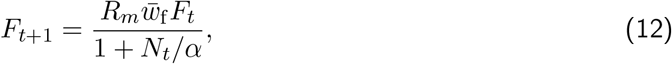

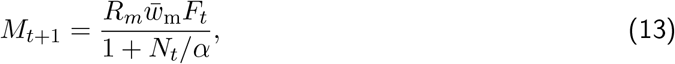

here *R*_*m*_ is the absolute mean number of offspring from females, and *α* is a parameter controlling density dependent growth, which we typically tune to give a carrying capacity 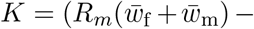 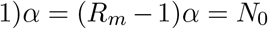, the initial population size we set for the population in the absence of drive. In these simulations we take a typical value of *R*_*m*_ = 6.

#### Multiplexed drive

In general, notational complexity increases considerably when we consider 2-fold and higher orders of multiplexed drive. We will only consider 2-fold and 3-fold multiplex in this paper, but we will set out a general framework for any order of multiplex *m*.

Firstly, how many different haplotypes *n*_*H*_ are there for each degree *m* of multiplex? For *n* = 3 alleles at each site (excluding drive), for 2-fold multiplex, as we calculated above there are *n*(*n* + 1)/2 = 6 haplotypes, which is the number of unordered pairs, corresponding to the number of upper diagonal elements for *n*×*n* matrix; including drive, we then need to track a total of *n*_*H*_ = 7 haplotype frequencies. For 3-fold multiplex, the number of haplotypes to be tracked is the number of unordered triplets, which is not so trivially calculated, but amounts to calculating the analogue of the number of “upper diagonal” elements of a *n×n×n* tensor. The answer is that there are *n*^(*m*)^/*m*! = 3^(3)^/3! = 10 haplotypes, excluding drive, for *m* = 3 fold multiplex, where *n*^(*m*)^ = *n*(*n* + 1)(*n* + 2)…(*n* + *m* 1) is the rising factorial — note this expression holds true for any *m* integer. So including drive, for 3-fold multiplex, there are a total of *n*_*H*_ = 11 haplotypes frequencies to be tracked.

We also need to decide on the ordering of the haplotypes in the frequency vectors ***x*** and ***y***. The convention we use is column ordering and its generalisation for higher dimensional tensors, which gives the mapping of indices to haplotypes as shown in Table 1 and Table 2

The fitness matrices for 2-fold multiplex are 7 × 7, while for 3-fold they are 11 × 11. For males their fitness matrices are filled with all ones. We do not explicitly write out the fitness matrix of female genotypes for multiplex drive, but its construction follows the same rules as for 1-fold drive described above, where there are 3 categories of genotypes, where now any haplotype with an N at any site becomes non-functional: 1) only functional haplotypes on both chromosomes, where fitness 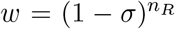, where *n*_*R*_ is number of R alleles across both chromosomes (e.g. genotype WR/RR has *n_R_* = 3, and *w*_23_ = *w*_32_ = (1 – *σ*)^3^); 2) functional/non-functional genotypes have fitness of the form 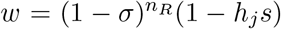 (e.g. genotype WRN/RRR has *n_R_* = 3 and *h*_*j*_ = *h*_*N*_, so *w*_46_ = *w*_64_ = (1 – *σ*)^3^ (1 – *h*_*N*_*s*), or genotype WWR/DDD has *n_R_* = 1 and *h*_*j*_ = *h*, so *w*_2,11_ = *w*_11,2_ = (1 – *σ*)(1 – *hs*)); 3) non-functional genotypes all have fitness *w* = 1 – *s* (e.g. *w*(WWN/DDD) = *w*_5,11_ = *w*_11,5_ = 1 – *s*).

The mutation matrix, for mutiplexed drive of degree *m*, ***M***^(*m*)^, is also a *n*_*H*_ × *n*_*H*_ matrix, the *j*^*th*^ column representing the rate of mutations from the *j*^*th*^ haplotype to each of the other haplotypes. We assume that the only mutations possible, at each target site, are W → R and W → N, with no backmutations. For this reason the only non-zero columns are haplotypes *j* that have at least a single W allele; of these each W alleles will mutate according the first column of the 1-fold mutation matrix, ***M*** (Eqn.8), whilst the alleles that are not wild type are unaffected by mutation and so are given by the second and third columns of ***M***. For mutation probability from the *j*^*th*^ haplotype to the *i*^*th*^, if we let *n*_*W*_ be the number of W alleles in the *j*^*th*^ haplotype, and 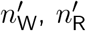 and 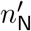 be the number of W, R, N alleles in the subset of the *i^th^* haplotype that only has W alleles, then for *m* = 2 the probability of mutation is

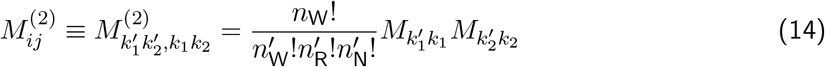

where haplotype *k*_1_*k*_2_ maps to *j* and haplotype 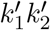 maps to *i*, using Table 1 and 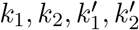 can take values {1, 2, 3} corresponding to alleles {W, R, N}, multinomial coefficient accounts for the redundancy in the number of ways haplotype 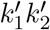. can be obtained by mutation of *k*_1_*k*_2_. Explicitly, evaluating this for *m* = 2 we arrive at the following mutation matrix

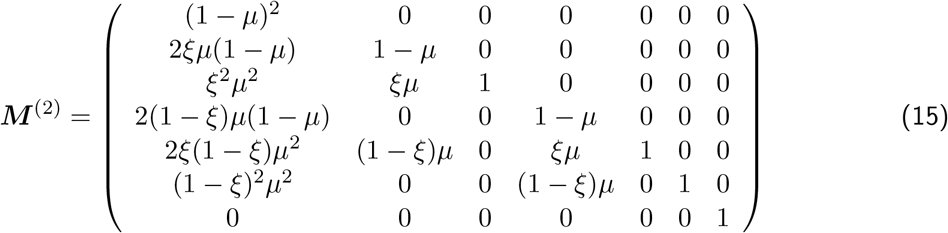

For *m* = 3, equivalent considerations give the following mutation probability from the *j*^*th*^ to *i^th^* haplotype

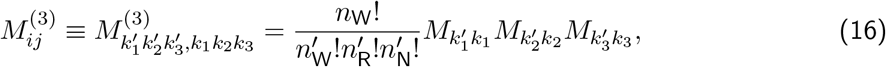

where haplotype *k*_1_*k*_2_*k*_3_ maps to *j* and haplotype 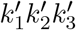 maps to *i*, using Table 2 and 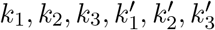 can take values {1, 2, 3} corresponding to alleles {W, R, N}, respectively.

For multiplex drive, we make the assumption that drive acts independently at each target site, as shown in Fig.2 for *m* = 2 gRNAs, and the genotype WW/DD, where we assume the same values of *ϵ*, *ν*, and *β* at each target site. The fraction of gametes is then effectively c alculated b y a multiplicative table of the fractions of W, R, N, D produced at each target site; since only a single successful cleavage and repair is needed (signified by a single D), we need to account for a ll fractions shown shaded in red in the table, to calculate the total fraction of DD gametes. This is formalised mathematically below and extended to any number of *m* of gNRAs.

To consider how multiplexed drive affects h a plotype f r equencies, w e n e ed t o c o nsider n ow that there is more than just a single genotype that is affected b y d r ive. A l l g enotypes w i th d r ive o n one chromosome and at least a single W allele on the other can be converted to homozygous drive; for *m* = 2 there are three which are enumerated in Table 3 and for *m* = 3 there are six enumerated in Table 4.

Each of these heterozygotes produces different fractions of the *n*_*H*_ possible haplotypes, depending on the number of W alleles in each haplotype. We create an analogous matrix to ***M*** (Eqn.8), whose *j*^*th*^ column gives the fraction of gametes produced by the *j*^*th*^ allele assuming it is in genotype with D:

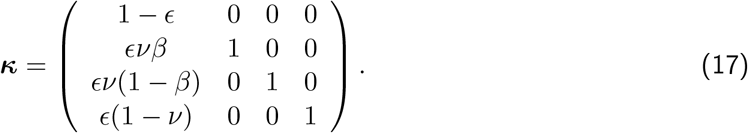

With this matrix the fraction of gametes that have the *i*^*th*^ haplotype from drive heterozygote *j* can be calculated for any value of *m*. For haplotypes *i* < *n*_*H*_ (i.e. not including drive) this is given by

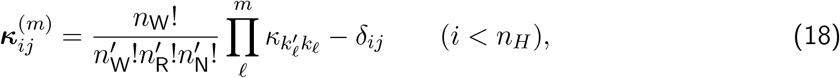

where *n*_W_ is the number of W alleles in the drive heterozygote *j*, and 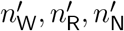 are the number of W, R, N alleles in haplotype *i*, and as before the haplotypes *k*_1_*k*_2_ and 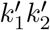 map to indices using Table 1. For *n*_*H*_, the drive haplotype (i.e. DD or DDD), we need to sum over all haplotypes *i* that produce at least a single D allele (which is effectively converted to, for example, DD or DDD); if these set of haplotypes is denoted by 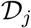 then

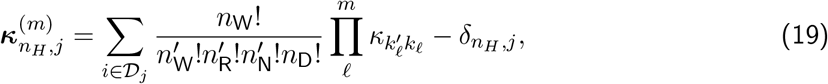

where on the RHS the relation between 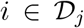 and 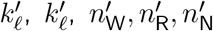 and 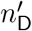 is implicit, and straightforward to enumerate.

Together these equations allow us to write down expressions for the update equations for haploype frequencies between generations for *m*-fold multiplex

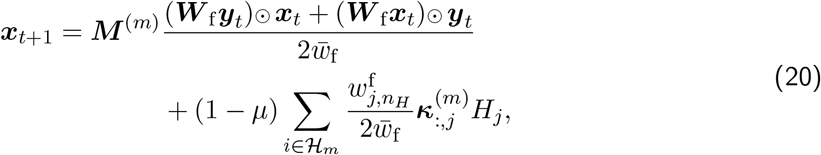

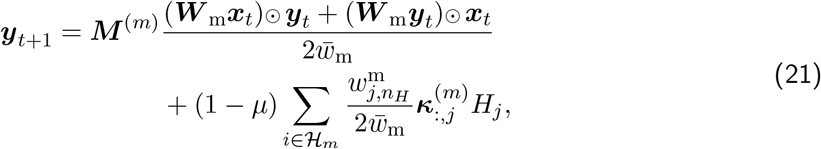

where 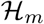 is the set of indices of haplotypes that have at least a single W allele, which is a function of *m* the degree of multiplexing (Tables 3&4), *H*_*j*_ are the drive-heterozygote frequencies given in Tables 3&4, where the time-dependence on *t* is implicit, and where the notation 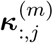 is the *j*^*th*^ column of the matrix ***κ***^(*m*)^.

#### Typical trajectories of allele/haplotype frequencies

In Fig.3 we show for *m* = 1, *m* = 2 and *m* = 3 the typical trajectories of the different alleles/haplotypes at population sizes for *N* < *N*^*^ (left hand plots of the figure), where resistance does not arise and *N* > *N*^*^ (right hand plots of the figure), where resistance does arise, due to de novo mutation or NHEJ. Below each is the corresponding time series of population size, showing population elimination (left) and rescue (right). These simulations assume that the fraction of functional NHEJ mutations is *β* = 10^−4^ and that all de novo SNPs are functional (*ξ* = 1). We see in all cases that initially drive replaces the wild type in less than 10 generations, which causes the mean population fitness to decrease and the population size itself to decrease; following this, if *N > N*^*^ we see that functional resistance mutants arise and then fix giving rise to population rescue, while for *N* < *N*^*^, resistance mutants do not arise on the timescale at which the population is eliminated. As we increase the degree of multiplexing *m*, the same story unfolds, except there are many more resistance haplotypes, which all arise at small frequency, and except for the single functional mutant for each value of *m* (R, RR and RRR, respectively for *m* = 1, 2, &3 gRNAs), none of these rise to sufficiently high frequency to achieve population rescue as they are deleterious as homozygotes, and as drive heterozygotes, haplotypes containing W can still be converted to D. The resistance haplotypes that have *m* copies of R do increase to high frequencies giving resistance and population rescue.

#### Concurrent vs Sequential accumulation of resistance alleles for NHEJ vs de novo mutation

When there are multiple gRNAs, whether by NHEJ or de novo SNPs, functional resistance mutations can arise sequentially (e.g. WWW → WWR → WRR → RRR) or concurrently through *m*-fold mutations (e.g. WWW → RRR) across the target sites, or some combination of the two (e.g. WWW → WWR → RRR). Here, using a simple heuristic analysis, we examine based on calculating the probability of fixation when there are a certain number of copies of functional resistance haplotypes, what signature we would expect in the scaling of *N*^*^ with respect to the base rate of generation of each type of mutant, which is *ϵβν* for NHEJ and *ξμ* for de novo SNPs. The approach essentially deterministically calculates the number of mutants generated over a certain timescale and ignores all fitness effects, assuming drift dominates at small frequencies, and only accounts for the fitness through the probability of establishment.

#### Generation of functional resistance non-homologous end joining mutants (NHEJ)

For a single gRNA, per generation and per each heterozygous W/D individual, non-homologous end joining mutants arise at a rate proportional to *ϵν*, and a fraction of *β* of these we assume to be functional (R), and so the rate of producing functional NHEJ mutants is proportional to *ϵνβ*. Hence, of *r*_*t*_ is the frequency of the R allele in the population in generation *t*, then the difference equation describing the dynamics is given by

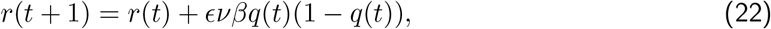

where *q*_*t*_ is the frequency of the D allele and to a good approximation *q*(*t*)(1 – *q*(*t*)) is the frequency of the W/D heterozygotes, assuming all other alleles apart from the W and D are at much smaller frequencies initially. Mutants generated earlier will generally have contribute most to the ultimate probability of establishment and population rescue, and so we make the approximation that the frequency of the heterozygotes is roughly *q*(*t*) and so

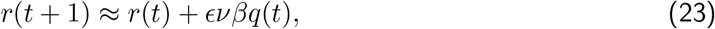

which has solution *r*(*t*) = *r*(0) + *ϵνβQ*(*t*) = *ϵνβQ*(*t*), since we assume at *t* = 0 when drive is introduced there are no mutants, and where 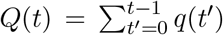. If we assume there is some characteristic time over which mutants are generated, related to the time to fixation of drive *τ*, then the number of copies of the mutant generated in this time is *τ* is therefore 2*Nr*(*τ*) = 2*NϵνβQ*(*03C4*). If each mutant has probability of fixation of *π* = 2*s*_*b*_, where *s*_*b*_ is an effective selection coefficient of the benefit the functional resistance allele has in the presence of drive, then given this number of mutants, we calculate the probability of fixation by 1- probability that none of these mutants establishes:

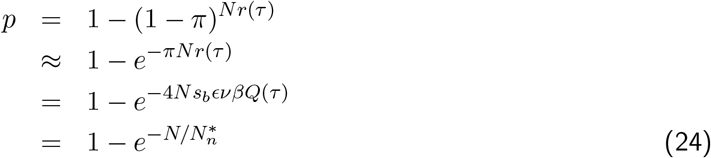

where 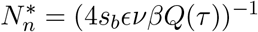, which gives us the result that we expect resistance to arise significantly, when *N* > *N*_*n*_ or the rate of generation of NHEJ mutants is 2*Nϵνβ* ~ (2*s*_*b*_*Q*(*τ*))^−1^, then the probability of resistance is large.

For *m* = 2 gRNAs, a single resistance mutation (WR or RW) is not sufficient to prevent the copying of drive and so only haplotypes with two functional resistance mutations (RR) have a selective advantage in the presence of drive. Assuming the frequency of all mutants is small, and so assuming the fact that single mutants are selected against can be ignored (due to drift), the difference equations describing this scenario are

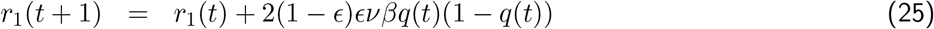

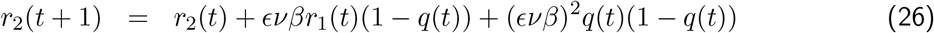

where *r*_1_ is the frequency of mutants with a single functional resistance allele at either of the two target sites and *r*_2_ is the frequency of mutants that have a functional resistance allele at both target sites. If we make the same assumption that initially the frequency of drive will be small, such that *q*(1 – *q*) ≈ *q*, then the solution for the frequency of single-fold functional resistance mutants is *r*_1_(*t*) = 2(1 – *ϵ*)*ϵνβQ*(*t*), and so we have a difference equation for only *r*_2_:

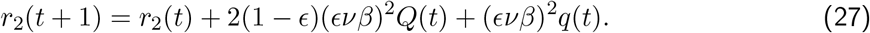

The second term represents the generation of two-fold mutants from single-fold heterozygotes with drive (WR/DD or RW/DD), whilst the third term represents generation directly from wild type-drive heterozygotes. We can see that if 1 – *ϵ* is small then later direct mechanism of generation will be dominant, as long as the cumulative frequency of drive *Q*(*t*) is not significantly larger than the current frequency *q*(*t*); as *Q*(*t*) is a cummulative frequency it will be of order 1/*s*_*D*_, where *s*_*D*_ = (1 + *ϵ*)(1 – *hs/*2) – 1 is the effective selection coefficient of drive, which gives the condition *s*_*D*_ – 2(1 – *ϵ*) such that direct generation of double mutants dominates over sequential generation. For typical parameters of suppression drive as used in this paper *ϵ* = 0.95, *s* = 1 and *h* = 0.3, we get *s*_*D*_ ≈ 0.66 and 2(1 – *ϵ*) = 0.1, and so we would expect direct generation of double mutants to dominate. If this is the case then our difference equation for double functional mutants is

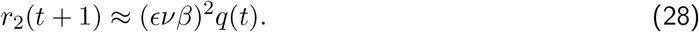

and so comparing to Eqn.22, we have the same difference equation but *ϵνβ* → (*ϵνβ*)^2^ and so would expect the probability of resistance to be of the form 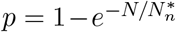 with 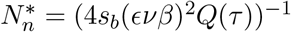. A similar analysis can be performed for *m* > 2 gRNAs to give in general 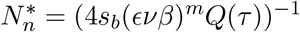, for *m* gRNAs. Here, we see that if we let *γ*_*n*_ = *s*_*b*_*Q*(*τ*) then we have a single unknown constant, which is the same for all *m*, and fitting to the simulation data of *p* vs *N* for different values of *β* (Fig.1 in main text), we find indeed this is the case and *γ* ≈ 0.2 for *m* = {1, 2, 3}, as shown in Fig.4.

#### De novo generation of SNP functional resistant mutants

In the case of de novo generation of mutants the difference equation for a single gRNA is

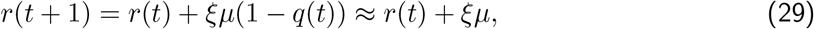

where we assume that initially *q* ≪ 1. The solution is simply *r*(*t*) = *ξμt*. Using the same simple heuristic as for NHEJ, this then leads to the probability of fixation of mutants generated up to some time *τ*, related to the timescale for fixation of drive, to be:

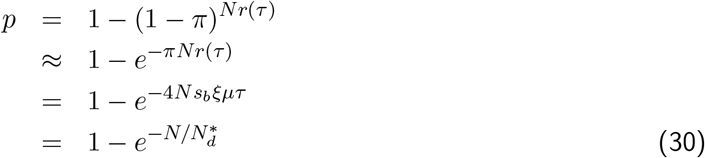

with 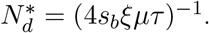

For *m* = 2 gRNAs, the difference equations for de novo generation of single and double functional resistance mutants is

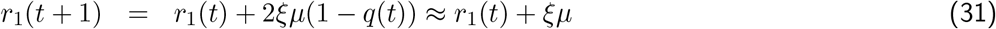

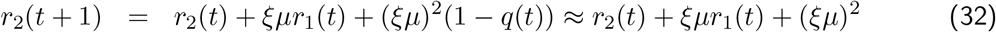

where again we assume the frequency of drive is initially small and we ignore any fitness effects for mutants at small frequency in calculating the frequency of mutants. As above the frequency of single mutants will be *r*_1_(*t*) ≈ 2*ξμt*, where now as there are two paths from the wild type to generate a single mutant there is a factor of 2, and so the difference equation for double functional mutants will be

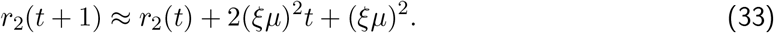

Now here the second term, representing the sequential generation of mutants, will dominate for any sufficiently nu mber of ge nerations *t* > 1 an d so ig noring the third term wh ich re presents concurrent generation of mutants, the solution is *r*_2_(*t*) = 2(*ξμ*)^2^*t*(*t* + 1)/2 ~ (*ξμ*)^2^*t*^2^ for large *t*. This means on a time scale *τ* the number of functional double-mutants generated will be 2*Nr*_2_(*τ*) ~ 2*N* ×(*ξμ*)^2^*t*^2^ = 2*N* (*ξμ*)^2^*t*^2^. Following the previous arguments, the probability of fixation will then be 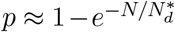 with *N*_*d*_ = (4(*ξμ*)^2^*s*_*b*_*τ*^2^)^−1^. Again the arguments can be extended to *m* = 3 gRNAs and we find *N*_*d*_ = (4(*ξμ*)^2^*s*_*b*_^3^)^−1^, to leading order in *τ*.

We can fit the simulation data of the de novo probability of resistance vs *N* as *ξ* is varied, using this heuristic theory with a single fitting constant *γ*_*m*_ and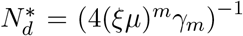, for each value of *m* and we find *γ*_1_ = 0.76 ± 0.004, *γ*_2_ = 6 ± 0.04 and *γ*_3_ = 55 ± 0.3, as shown in Fig.5; using the relation *γ*_*m*_ = *s*_*b*_*τ*^*m*^, we find these data are consistent with *τ* ≈ 8.5 generations and a beneficial selection coefficient of *s*_*b*_ ≈ 0.09.

#### Multiplex resistance from standing variation of SNP mutants

In this section, we show that the significant amplification of standing variation or pre-existing mutants for *m* > 1 gRNAs can be heuristically understood by calculating approximately the average probability of fixation of *m*-fold functionally resistant, assuming that in the presence of drive it has a beneficial selection coefficient *s*_*b*_ and that before the introduction of drive it is mildly selected against, giving a distribution of the frequency of such mutants in mutation-selection balance.

For *m* = 1 gRNAs the relevant theory has already been calculated in detail by Hermisson & Pennings (*2*), which we reproduce here in brevity and slightly simplified form. In mutation-selection balance, the diffusion approximation can be used to calculate the equilibrium distribution of the frequency of such variants. Here a single functional resistance mutant has selection coefficient –*σ* before the introduction of drive and the distribution of frequency *r* is approximately gamma distributed, as long as *α* = 4*Nσ* ≫ 1:

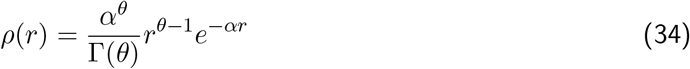

which has shape parameter *θ* = 4*Nξμ* and rate *α*. The mean of this distribution is 〈*r*〉 = *θ/α* = *ξμ/σ*, which is the classic mutation-selection balance frequency. Now the probability of fixation given an initial frequency *r*_0_ and beneficial selection coefficient *s*_*b*_ is

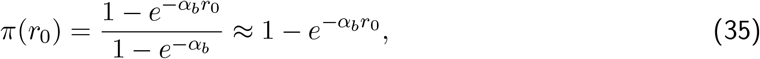

where the last approximation assumes *α*_*b*_ = 4*Ns*_*b*_ – 1. The average probability of fixation assuming a distribution *ρ*(*r*_0_) is then simply calculated by averaging over the initial frequency *r*_0_, which gives

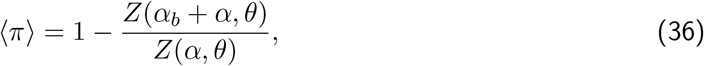

where *Z* is the normalisation constant of the gamma distribution with population scaled fitness *f* : 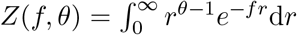, where extending the integral to ∞ will have small penalty in the strong selection limit, i.e. as long as *f* ≫ 1. Evaluating this and re-arranging we find the result of Hermisson & Pennings in the strong selection limit:

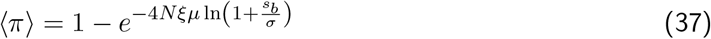

And so the probability of resistance 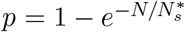, where 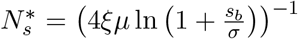, which is of the same form as for NHEJ and de novo SNP mutations. There is only a single unknown parameter *s*_*b*_, and so fitting this form to the simulation data for the *p* vs *N*, for varying *ξ* we find *s*_*b*_ ≈ 0.5 as shown in Fig. (*m* = 1 panel), which is larger than found from de novo simulations, and likely reflects some mechanistic aspect not reflected in these simple heuristic considerations.

For *m* = 2, we could try to calculate exactly the joint steady-state distribution of the frequency of all mutants, under the sequential mutation scheme WW → {WR, RW} → RR, but this is in general an unsolved problem in population genetics, since there is a net flux or current of probability through the states of the system (detailed balance is not obeyed) and so calculating this joint distribution is not straightforward. Here, instead we appeal to a simple heuristic, which reproduces the scaling with *σ* seen in the simulation data; we assume the distribution of the double-mutant frequency *r*_2_ follows the same gamma distribution, but that mutation to this allele is solely from single mutants whose frequency is assumed fixed at it's equilibrium frequency 〈*r*_1_〉 = 2*ξμ/σ*, where the factor 2 arises from the multiplicity 2 of paths to single mutants from the wild type; this is reasonable given that sequential mutation rather than concurrent is dominant, as demonstrated for de novo mutation. Given this and that the fitness cost of double mutants, before drive is introduced, is −2*σ*, the distribution of the double functional resistance mutants is

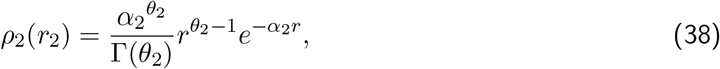

where *α*_2_ = 8*Nσ*, and *θ*_2_ = 4*Nξμ 〈r〉*_1_ = 8*N* (*ξμ*)^2^/*σ*. The average frequency of double functional mutants is *θ*_2_/*α*_2_ = (*ξμ*)^2^/*σ*^2^, which is simple to show is what we would expect solving the deterministic equations for the frequency of double-mutants. Using the form of Eqn.36, we find

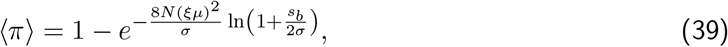

which is of the form 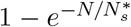 with 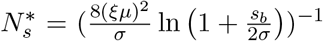, which we see means 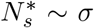 for *m* = 2, instead of the much weaker logarithmic dependence for *m* = 1. A similar analysis can be done for *m* = 3, which gives 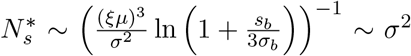, which is consistent with the power law sca ng we see in the simu tions (Fig.3b). In practice, we find that the following form 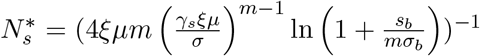 fits the simulation data of *p* vs *N* for different values of *ξ* and *m* with a single fitting parameter *γ*_*s*_ ≈ 2 (for *m* = 2 and *m* = 3) as shown in Fig.6.

Note that there is an implicit assumption in all these calculations, that establishment must arise before the population is removed, which will be true if ~ 1/*s*_*b*_ is much smaller than the timescale over which the population is removed.

#### Time to Resistance

In this section we plot the mean time to resistance over *M* = 500 replicates. Figs.7,8 & 9 are simulations for NHEJ only, Figs.10,11 & 12 are simulations for de novo only, and Figs.13,14 & 15 are simulations for preexisting and de novo only.

---

**Figure 1:**
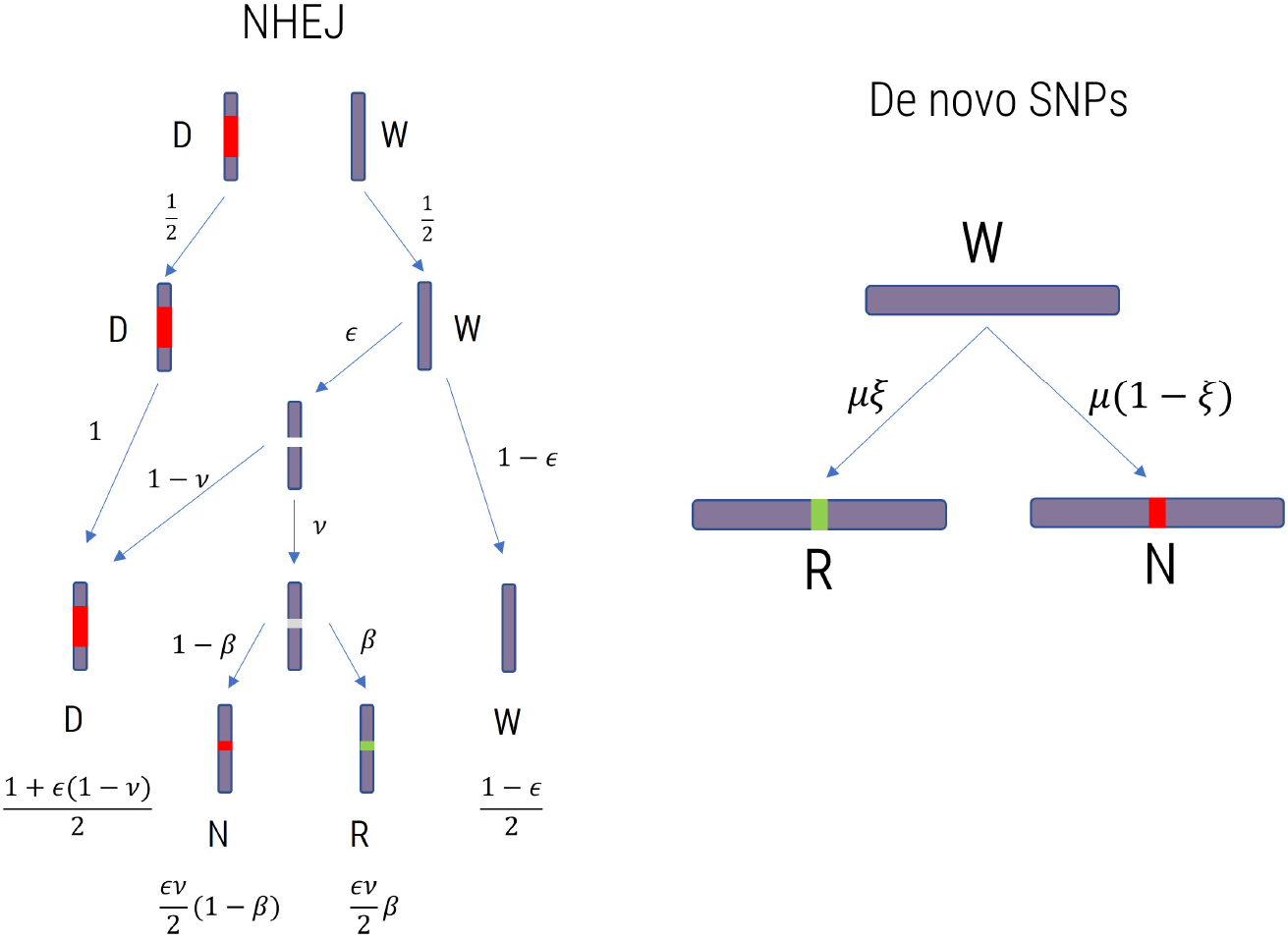
Diagram showing gametogenesis for *m* = 1 gRNAs with NHEJ for a W/D individual (left) and how de novo SNPs arise by mutation (right). On the right, wild type is the background colour, a white marker indicates the cleavage state, a long red bar represents drive inserted at the target site, a grey marker is an NHEJ event, which are functional (green) or non-functional (red).

**Figure 2:**
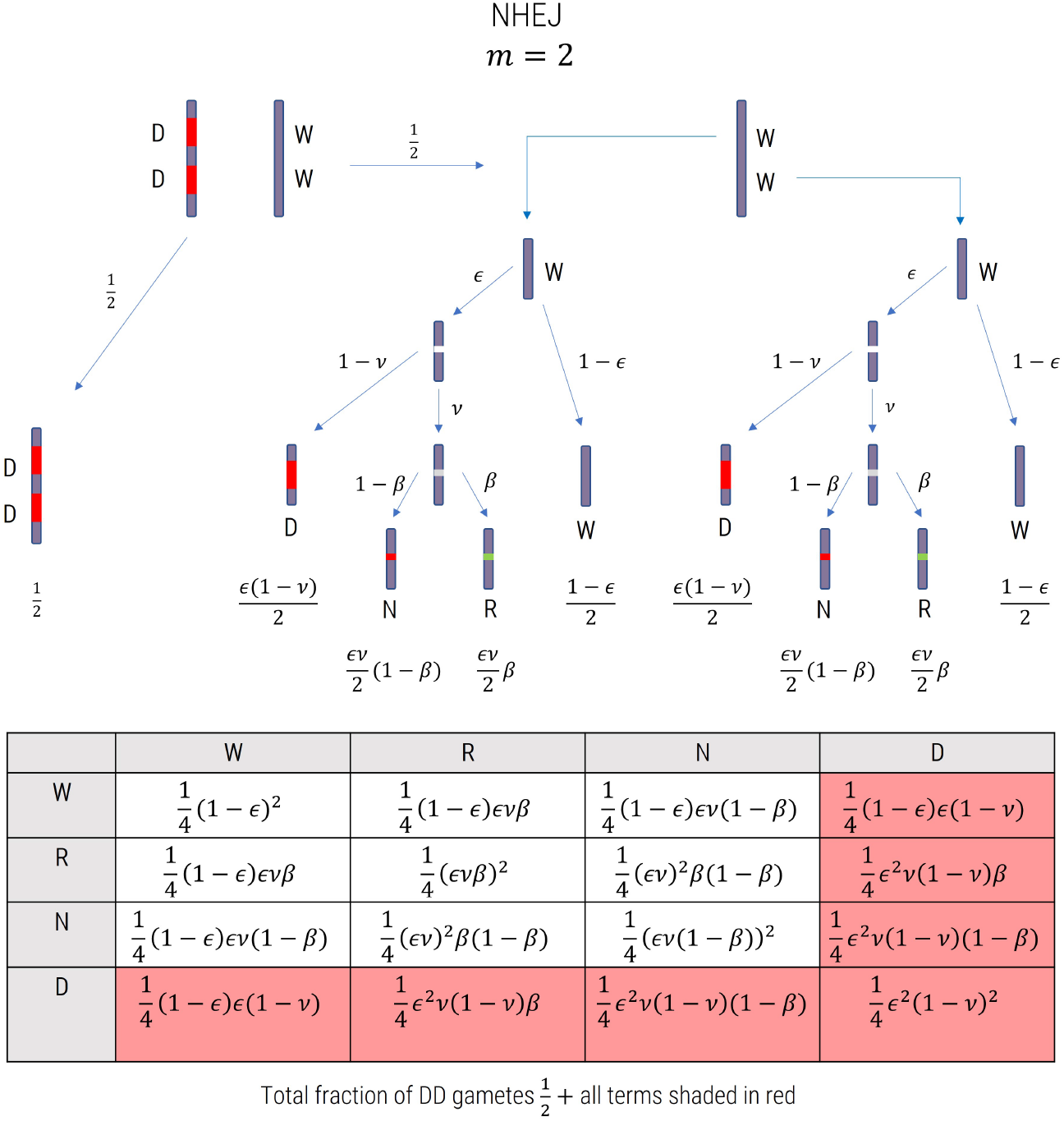
Diagram showing gametogenesis for *m* = 2 gRNAs with NHEJ for a WW/DD individual, where the model assumes the action of drive is independent on each target site. wild type is the background colour, a white marker indicates the cleavage state, a long red bar represents drive inserted at the target site, a grey marker is an NHEJ event, which are functional (green) or non-functional (red).

**Figure 3:**
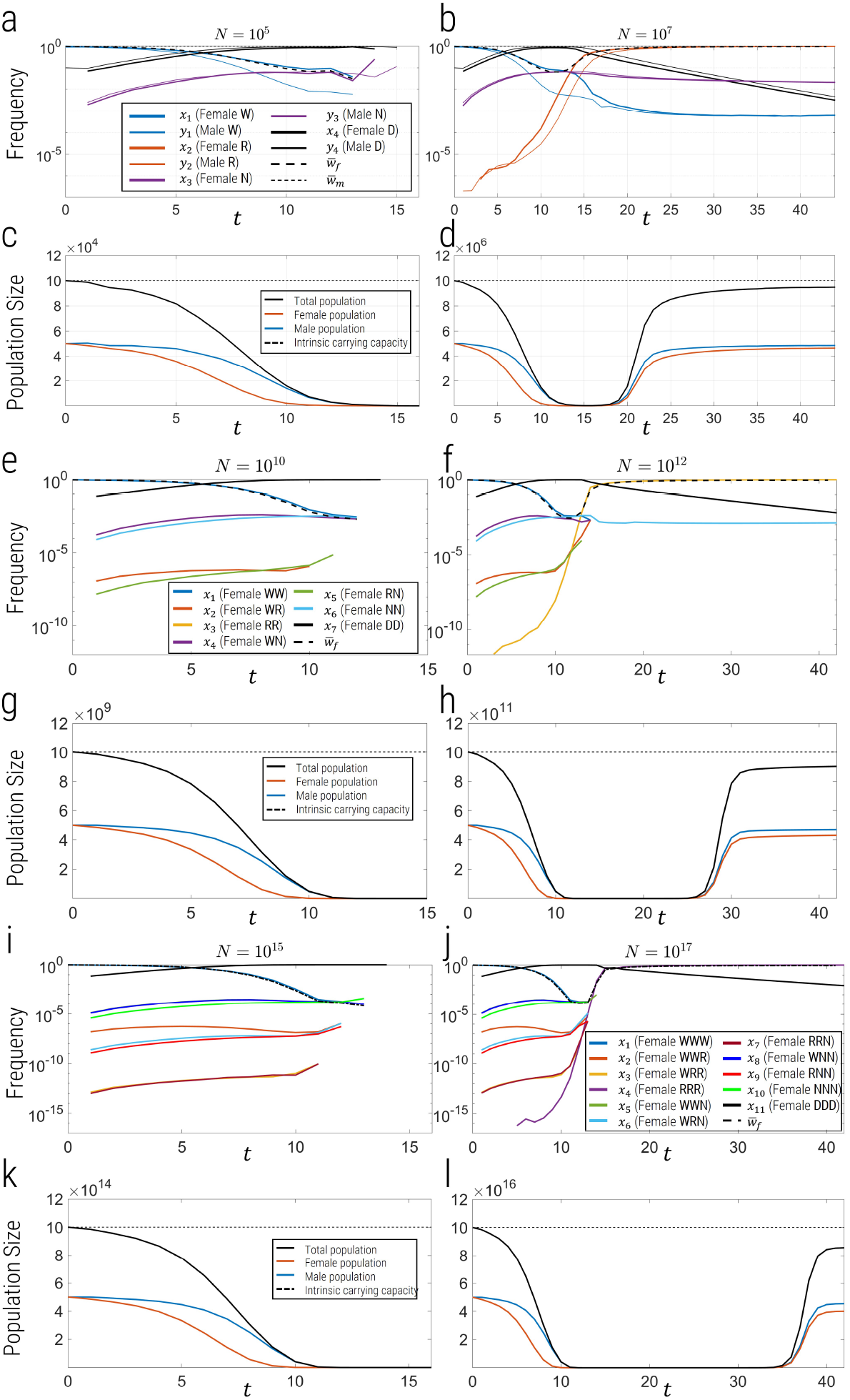
Plot of frequency of different alleles (a,b, e, f, i & j) and population dynamics (c, d, g, h, k & l) vs generation time for *N* < *N*^*^ (a, c, e, g, i & k) and *N* > *N*^*^ (b, d, h, j & l) for *m* = 1 gRNA (top panel), *m* = 2 gRNAs (middle panel) and *m* = 3 gRNAs (bottom panel), where *N*^*^ is the critical population size at which the probability of resistance (rescue) is 1 – *e*^−1^. For *m* = {2, 3}, we only plot the female allele frequencies for clarity.

**Figure 4:**
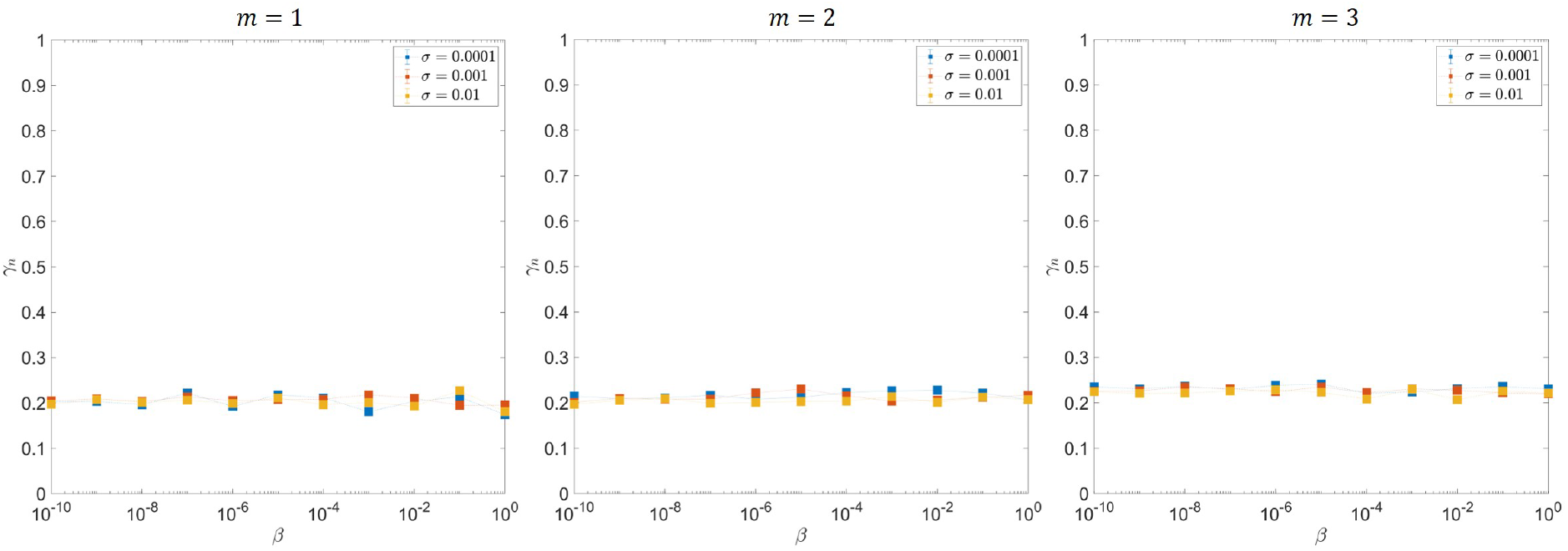
Values of *γ*_*n*_ fitting parameter from fits to simulation results of probability of resistance *p* vs *N* for different values of *β*, where 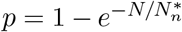 with 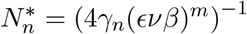, where each panel are for different numbers of gRNAs *m* as indicated.

**Figure 5:**
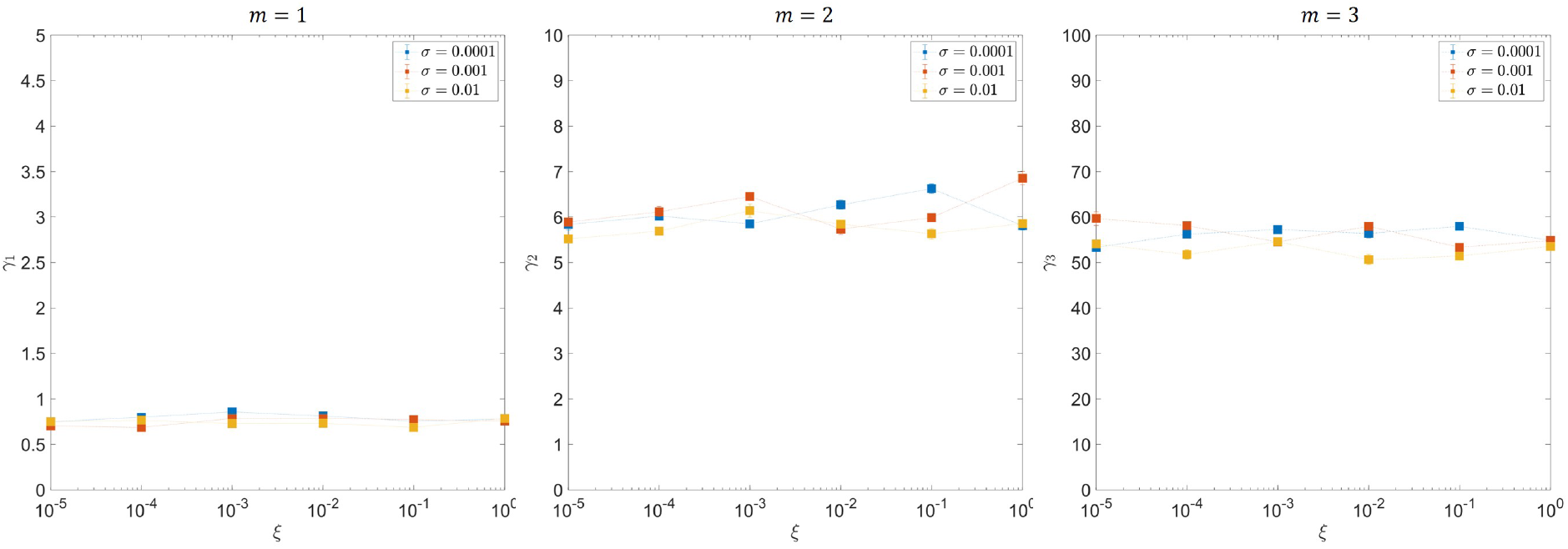
Values of *γ*_*m*_ fitting parameter from fits to simulation results of probability of resistance *p* vs *N* for different values of *α*, where 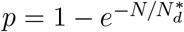 with 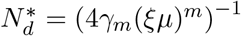, where each panel are for different numbers of gRNAs *m* as indicated.

**Figure 6:**
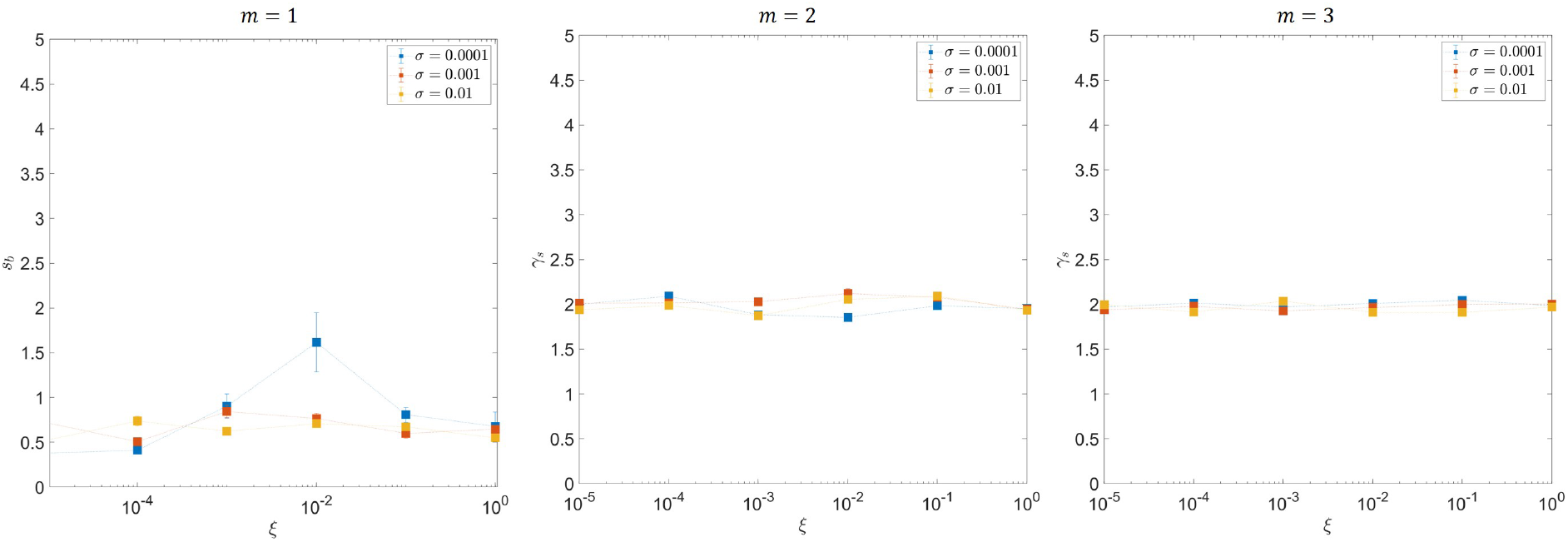
Values of *s*_*b*_ (*m* = 1) and *γ*_*s*_ (*m* = 2 and *m* = 3) fitting parameter from fits to simulation results of probability of resistance *p* vs *N* for different values of *ξ* with standing variation and de novo SNPs and NHEJ turned off, where 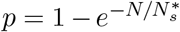 with 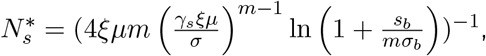 where each panel are for different numbers of gRNAs *m* as indicated.

**Figure 7:**
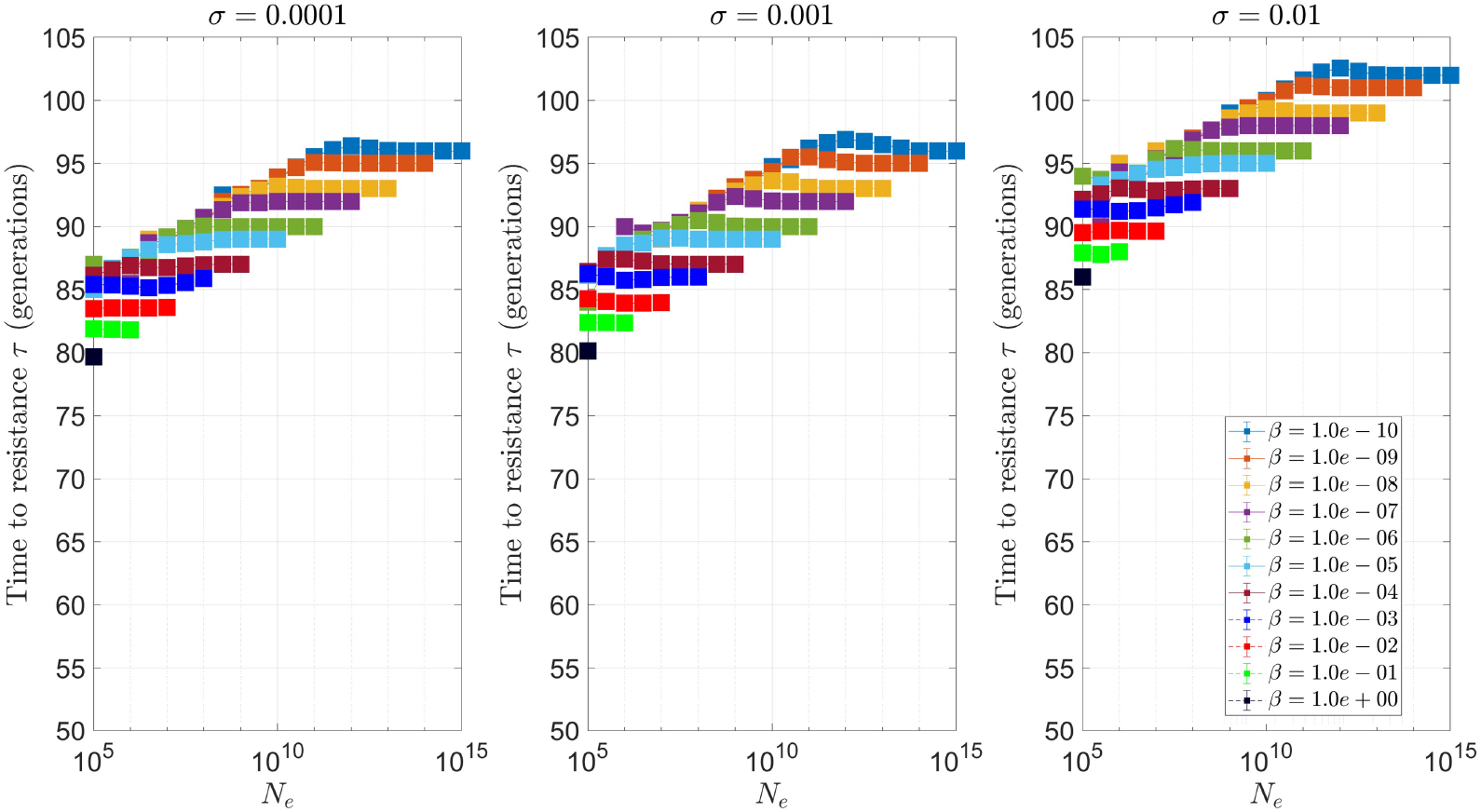
NHEJ only. Mean time to resistance over *M* = 500 replicate simulations for *m* = 1 gRNA. Standard parameters of simulations.

**Figure 8:**
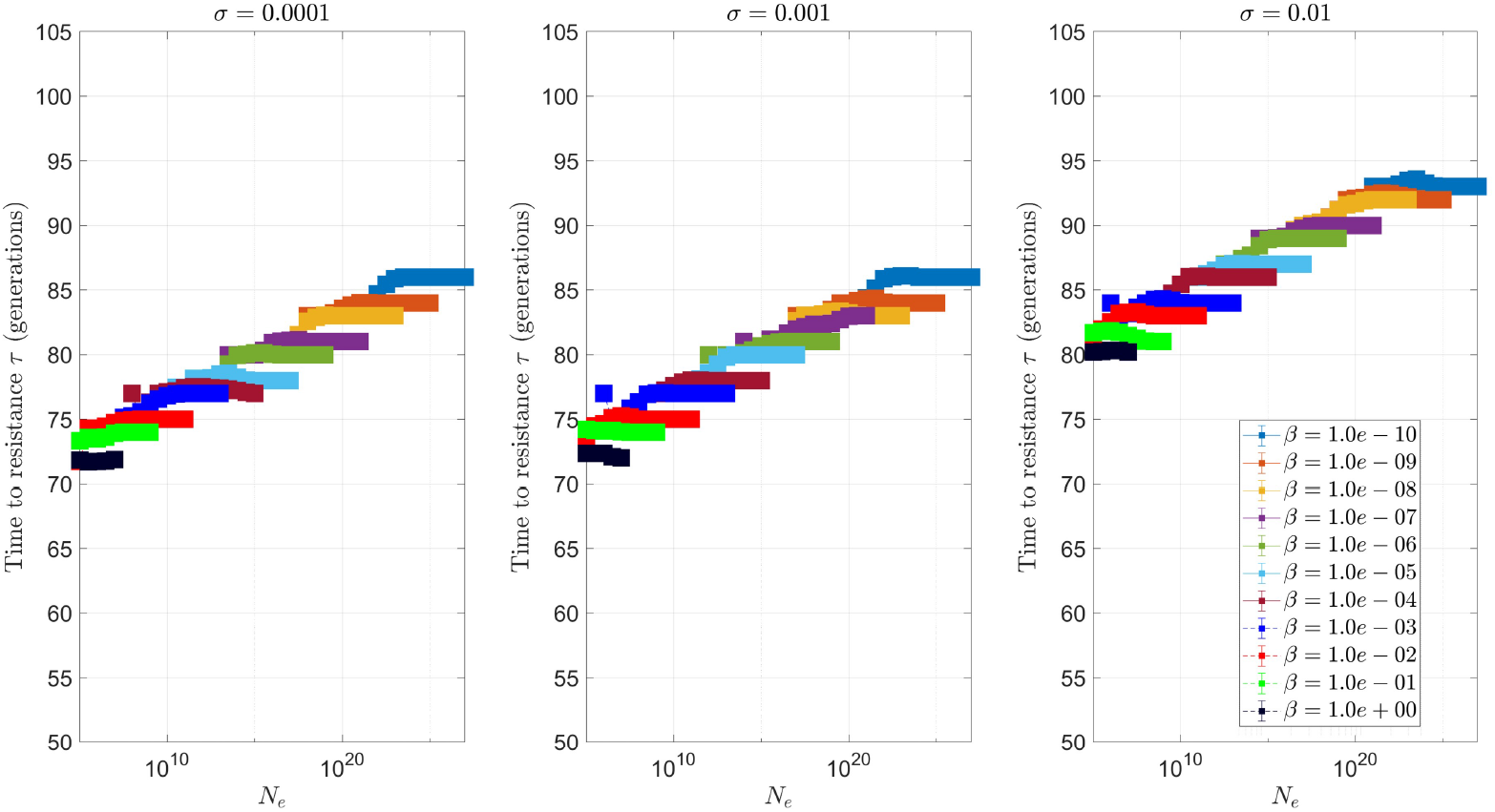
NHEJ only. Mean time to resistance over *M* = 500 replicate simulations for *m* = 2 gRNAs. Standard parameters of simulations.

**Figure 9:**
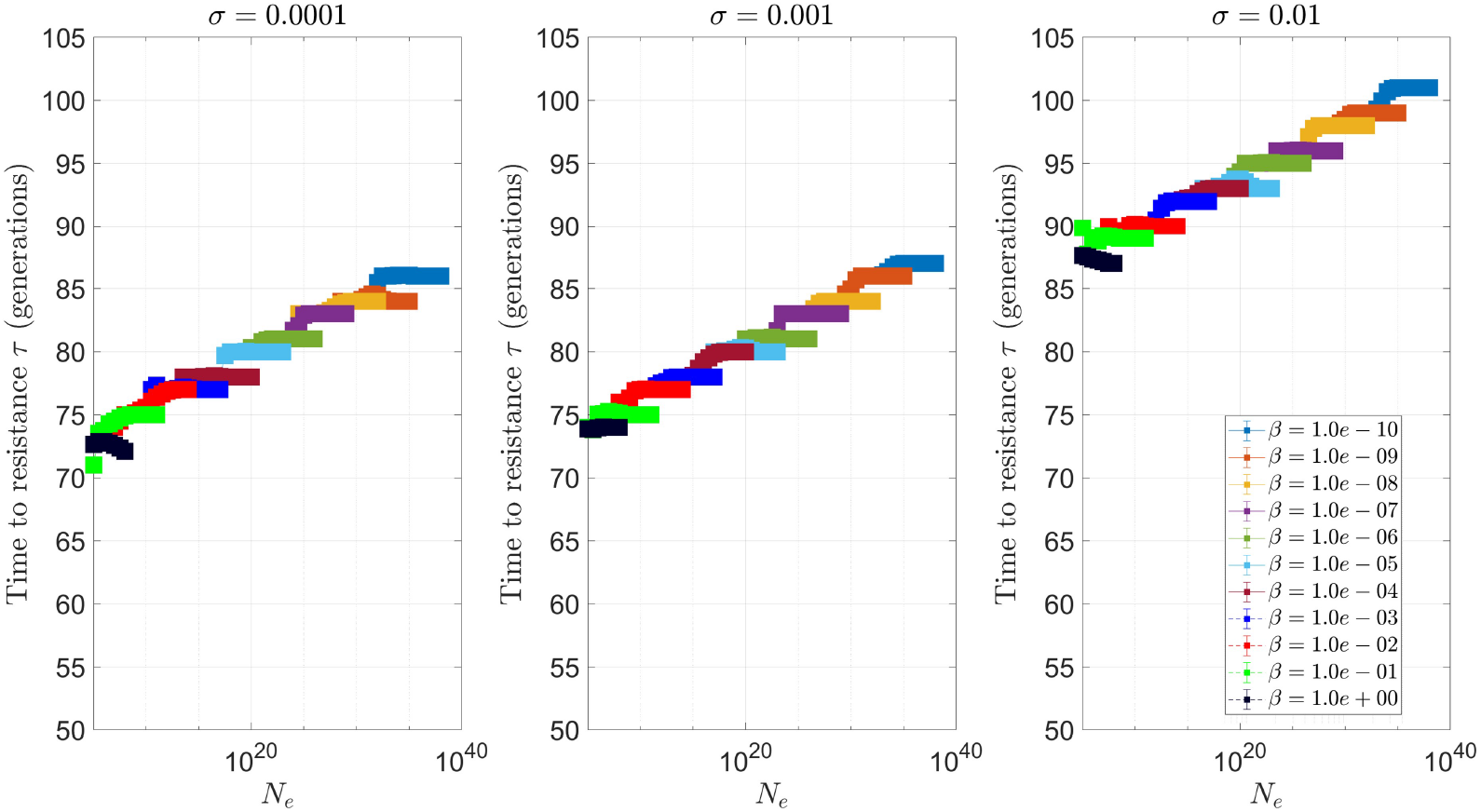
NHEJ only. Mean time to resistance over *M* = 500 replicate simulations for *m* = 3 gRNAs. Standard parameters of simulations.

**Figure 10:**
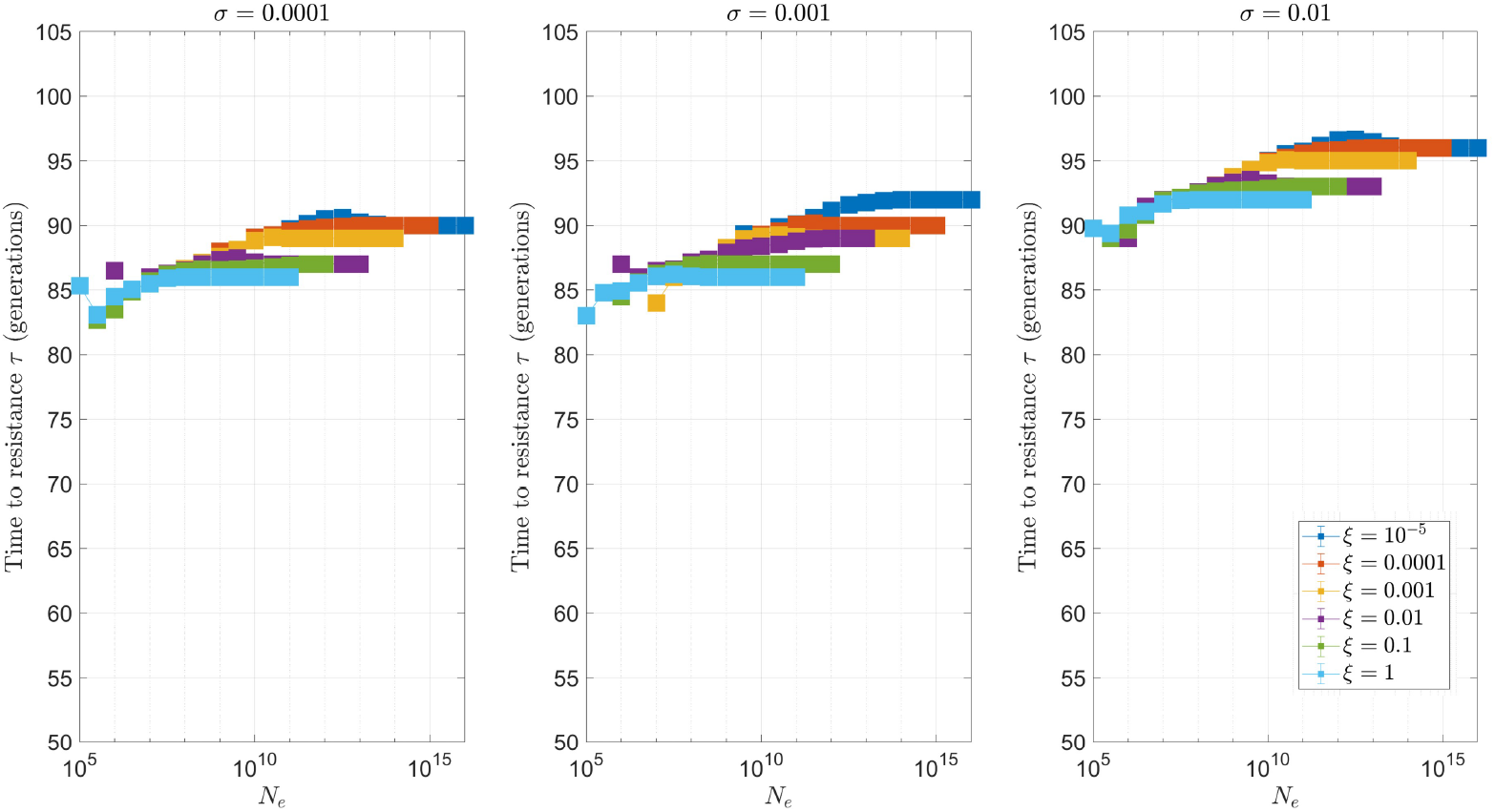
Single nucleotide mutations only (de novo). Mean time to resistance over *M* = 500 replicate simulations for *m* = 1 gRNA. Standard parameters of simulations.

**Figure 11:**
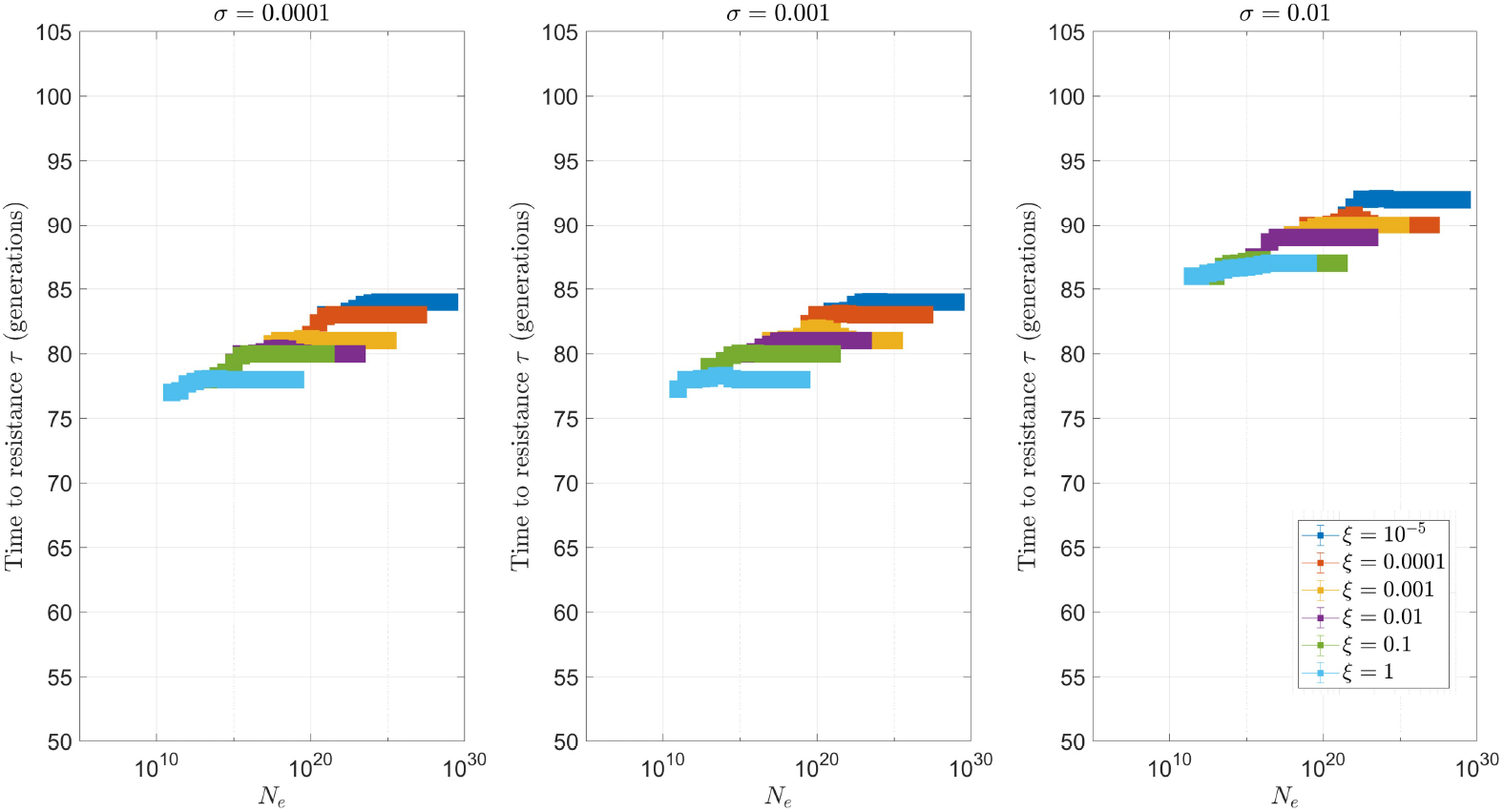
Single nucleotide mutations only (de novo). Mean time to resistance over *M* = 500 replicate simulations for *m* = 2 gRNAs. Standard parameters of simulations.

**Figure 12:**
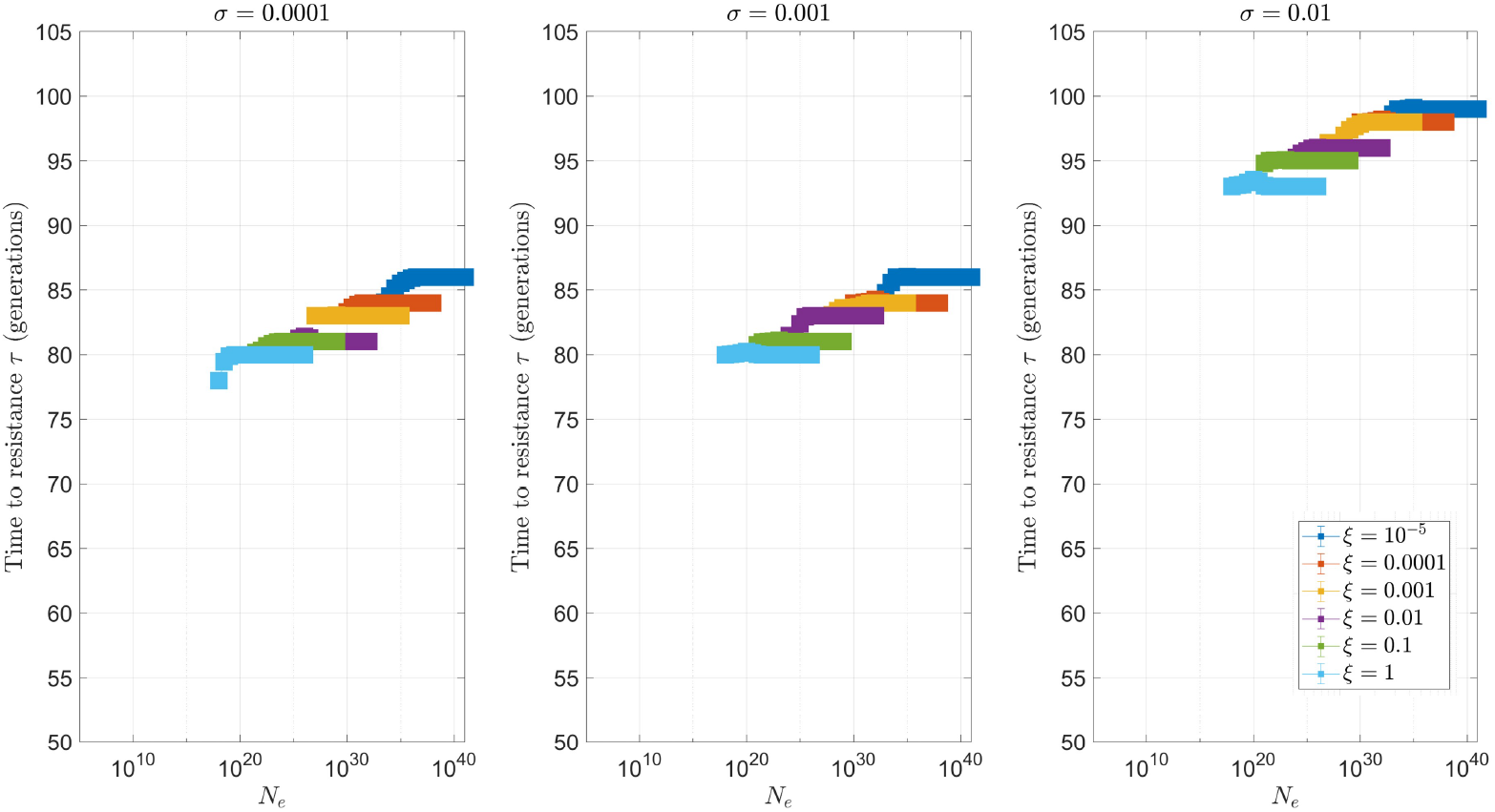
Single nucleotide mutations only (de novo). Mean time to resistance over *M* = 500 replicate simulations for *m* = 3 gRNAs. Standard parameters of simulations.

**Figure 13:**
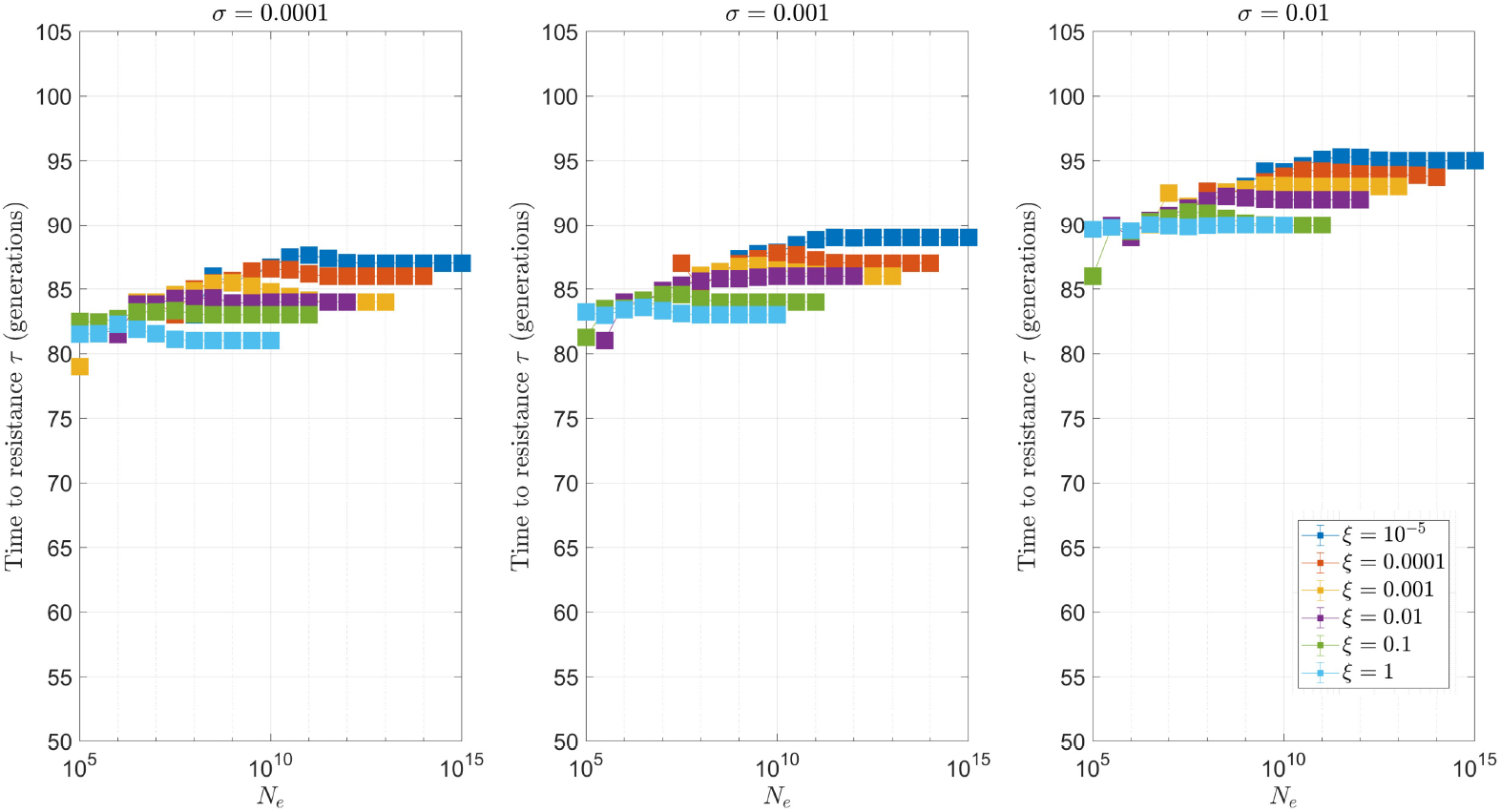
Single nucleotide mutations only (pre-existing + de novo). Mean time to resistance over *M* = 500 replicate simulations for *m* = 1 gRNA. Standard parameters of simulations.

**Figure 14:**
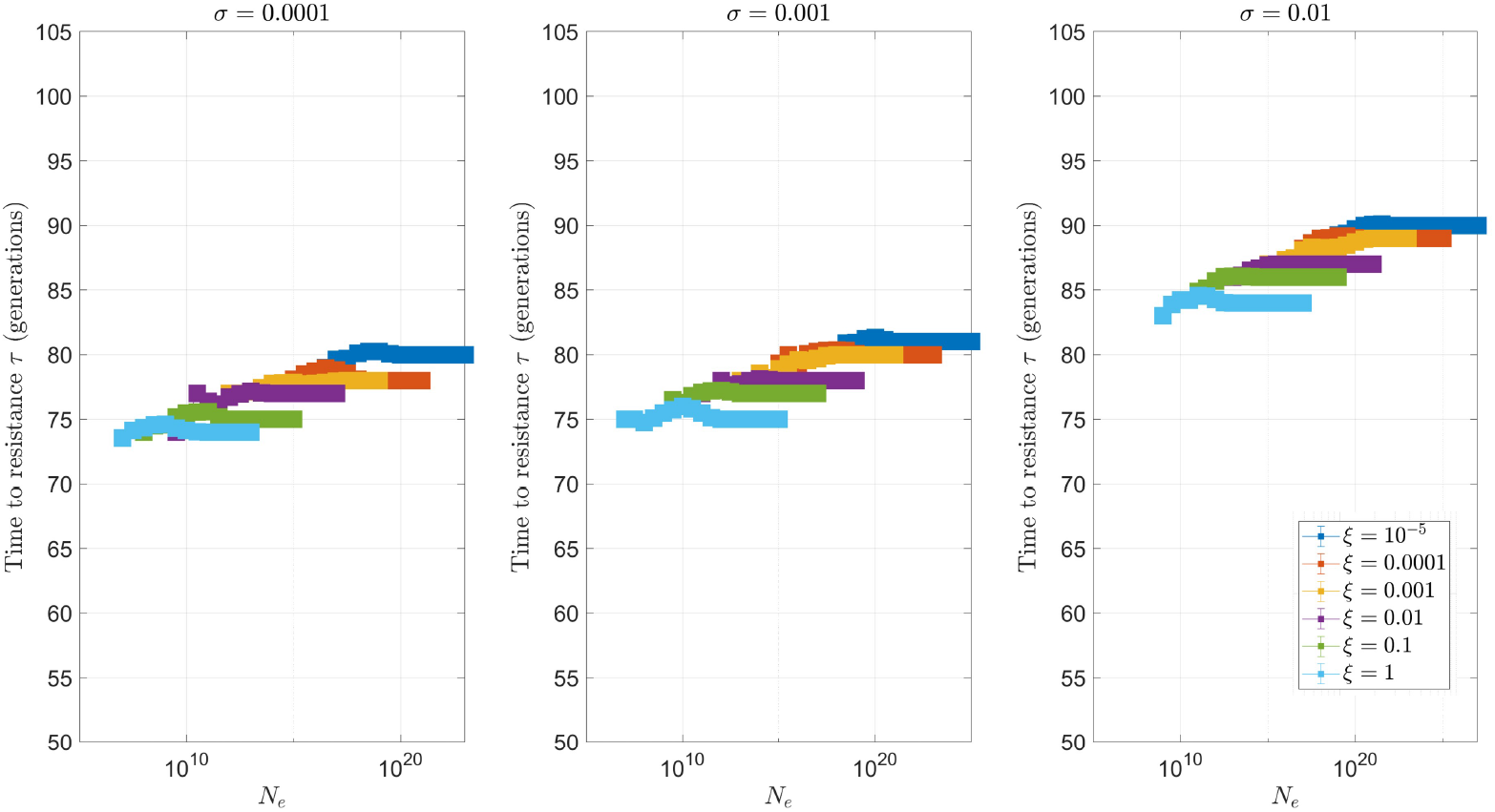
Single nucleotide mutations only (pre-existing + de novo). Mean time to resistance over *M* = 500 replicate simulations for *m* = 2 gRNAs. Standard parameters of simulations.

**Figure 15:**
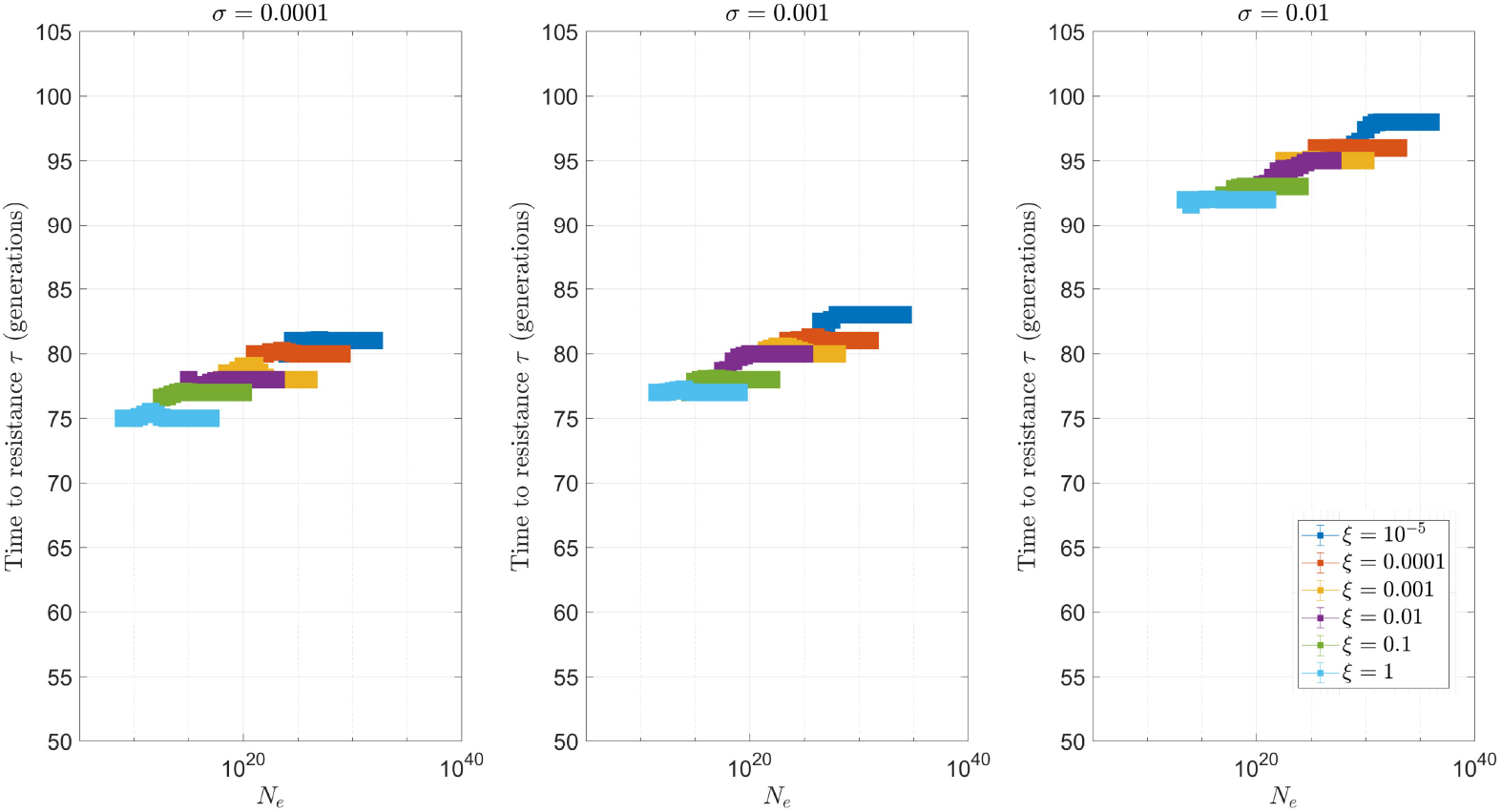
Single nucleotide mutations only (pre-existing + de novo). Mean time to resistance over *M* = 500 replicate simulations for *m* = 3 gRNAs. Standard parameters of simulations.

**Table 1:**
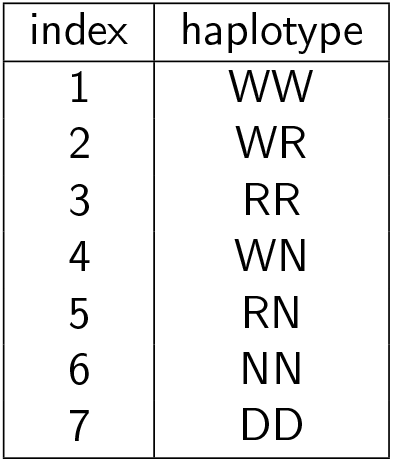
Mapping between frequency vector index and haplotypes for 2-fold multiplex

**Table 2:** Mapping between frequency vector index and haplotypes for 3-fold multiplex

| index | haplotype |
| --- | --- |
| 1 | WWW |
| 2 | WWR |
| 3 | WRR |
| 4 | RRR |
| 5 | WWN |
| 6 | WRN |
| 7 | RRN |
| 8 | WNN |
| 9 | RNN |
| 10 | NNN |
| 11 | DDD |

**Table 3:** Drive heterozygotes for 2-fold multiplex and their frequencies. Time dependence is implicit.

| genotype | heterozygote frequency |
| --- | --- |
| WW/DD | $H_1 = x_1 y_7 + x_7 y_1$ |
| WR/DD | $H_2 = x_2 y_7 + x_7 y_2$ |
| WN/DD | $H_4 = x_4 y_7 + x_7 y_4$ |

**Table 4:** Drive heterozygotes for 3-fold multiplex and their frequencies. Time dependence is implicit.

| genotype | heterozygote frequency |
| --- | --- |
| WWW/DDD | $H_1 = x_1 y_{11} + x_{11} y_1$ |
| WWR/DDD | $H_2 = x_2 y_{11} + x_{11} y_2$ |
| WRR/DDD | $H_3 = x_3 y_{11} + x_{11} y_3$ |
| WWN/DDD | $H_5 = x_5 y_{11} + x_{11} y_5$ |
| WRN/DDD | $H_6 = x_6 y_{11} + x_{11} y_6$ |
| WNN/DDD | $H_8 = x_8 y_{11} + x_{11} y_8$ |

## Notes

### Competing Interest Statement

The authors have declared no competing interest.

### Summary of Updates

Edited down for submission. Otherwise small changes.

